# DNA-triggered AIM2 condensation orchestrates immune activation and regulation

**DOI:** 10.64898/2026.03.18.712789

**Authors:** Quanjin Li, Xiaohan Geng, Huiwen Yan, Zhaolong Li, Miao Shi, Ziqi Zhu, Tongxin Niu, Chunqiu Zhao, Kaile Shu, Yina Gao, Han Feng, Songqing Liu, Qiuyao Jiang, Pengcheng Bu, Dong Li, Pu Gao

## Abstract

The innate immune sensor AIM2 detects cytosolic DNA and initiates inflammatory responses, yet its activation mechanism remains incompletely understood. Here, we show that AIM2 undergoes liquid-liquid phase separation upon DNA binding, forming dynamic condensates both *in vitro* and in cells. These condensates serve as platforms for inflammasome and PANoptosome assembly, promoting immune activation across multiple pathways. Direct structural determination from condensates reveals the assembly of active-form ASC filaments. Mechanistically, liquid-phase condensation is governed by multivalent interactions involving different AIM2 domains, including previously uncharacterized regions and species-specific elements. *In vitro* and *in vivo* assays show that mutants specifically disrupting condensation impair immune complex assembly, cell death initiation, antimicrobial defense, and intestinal homeostasis. Moreover, AIM2-DNA condensates function as regulatory hubs targeted by host- and pathogen-derived factors to balance immune homeostasis or facilitate immune evasion. These findings establish liquid-phase condensation as a fundamental mechanism of AIM2 activation and a potential therapeutic target.

## Introduction

Aberrant DNA resulting from pathogen infection or cellular stress serves as a critical pathogen- or damage-associated molecular pattern (PAMP or DAMP), which is detected by pattern recognition receptors (PRRs) to initiate innate immune responses. Among these PRRs, absent in melanoma 2 (AIM2) plays a crucial role in recognizing cytosolic double-stranded DNA (dsDNA) and recruiting the adaptor ASC and protease caspase-1 to form inflammasome.^1–5^ Inflammasome assembly induces autoprocessing of caspase-1, which subsequently cleaves downstream substrates gasdermin D (GSDMD) and proinflammatory cytokines pro-IL-1β and pro-IL-18, driving pyroptosis and inflammation.^6–14^ Beyond canonical inflammasome signaling, AIM2 also assembles with proteins such as ZBP1, RIPK1, RIPK3, FADD, and caspase-8 to form the AIM2 PANoptosome, a multi-protein complex that mediates inflammatory cell death (PANoptosis) by integrating pyroptotic, apoptotic, and necroptotic pathways.^15–18^ AIM2-mediated immune responses play vital roles in infection defense, tumor suppression, and neurodevelopment.^19–27^ However, its aberrant activation is associated with autoimmune and cardiovascular diseases, such as systemic lupus erythematosus and atherosclerosis.^22,24,28–31^ Thus, precise regulation of AIM2 is essential for maintaining immune homeostasis. Host proteins like p202 act as negative regulators of AIM2 inflammasome;^4,32,33^ while pathogens, such as HSV-1, encode inhibitors like VP22 to suppress AIM2 activation and evade immune detection.^34,35^

AIM2 comprises an N-terminal pyrin domain (PYD) for homotypic oligomerization, a C-terminal HIN domain for DNA binding, and a flexible intrinsically disordered region (IDR) connecting them. *In vitro* structural and biochemical studies suggest a potential activation model where multiple AIM2 molecules bind to the same dsDNA strand via their HIN domains, with peripheral PYD domains oligomerizing into a filamentous nucleation site that promotes ASC assembly and inflammasome formation.^5,36–41^ However, current understanding does not fully explain many observations and functional aspects of AIM2 activation. It is unclear how relatively low levels of AIM2 efficiently and simultaneously converge on a focused dsDNA region within the vast cytosolic space. The formation of the nucleation site requires the assembly of adjacent PYDs, while their corresponding HIN domains occupy only ∼20 base pairs (bp) of DNA, which is far shorter than the >80 bp needed for minimal AIM2 activation and the >300 bp required for optimal activation.^36,38,40,41^ While the model focuses on HIN-DNA and PYD-PYD interactions, cancer-associated mutations in the IDR and other “non-essential” regions imply that these areas may also contribute to AIM2 function (Figure S1A). Furthermore, the current simplistic interaction model does not explain how AIM2 coordinates the recruitment of multiple PANoptosome components and how its activity is regulated by various host- and pathogen-derived factors. These discrepancies highlight important gaps in our understanding of AIM2 activation and regulation, underscoring the necessity for additional in-depth mechanistic insights.

## RESULTS

### DNA induces AIM2 to form liquid-phase condensation *in vitro* and in cells

Upon stimulation with DNA or pathogen, endogenous mouse (m) and human (h) AIM2 robustly form puncta with DNA and ASC in mouse bone marrow-derived macrophages (BMDMs) and human monocytic THP-1 cells (Figures S1B and S1C). We reasoned that elucidating the characteristics and assembly mechanisms of AIM2 puncta could provide critical insights into AIM2-mediated signaling activation. To explore the properties of AIM2 puncta *in vitro*, we expressed and purified full-length mAIM2 and hAIM2 with an MBP tag. Upon mixing with 100-bp dsDNA and treatment with TEV protease to remove the MBP tag, mAIM2 rapidly forms liquid-like droplets with dsDNA (Figures 1A and S1D). As time progressed, mAIM2-DNA droplets fuse with each other into larger ones, accompanied by increased droplet size and fluorescence intensity (Figures 1A-1D). Fluorescence recovery after photobleaching (FRAP) experiments show efficient fluorescence recovery of mAIM2-DNA droplets after bleaching (Figures 1E and S1E), demonstrating their dynamic nature and efficient molecular exchange with the external environment—a hallmark of typical liquid-liquid phase separation (LLPS). Consistent with mAIM2, hAIM2 is also induced by dsDNA to form prominent condensates *in vitro*, but with a stronger propensity, exhibiting gel-like condensation characteristics (Figure S1F).

**Figure 1.**
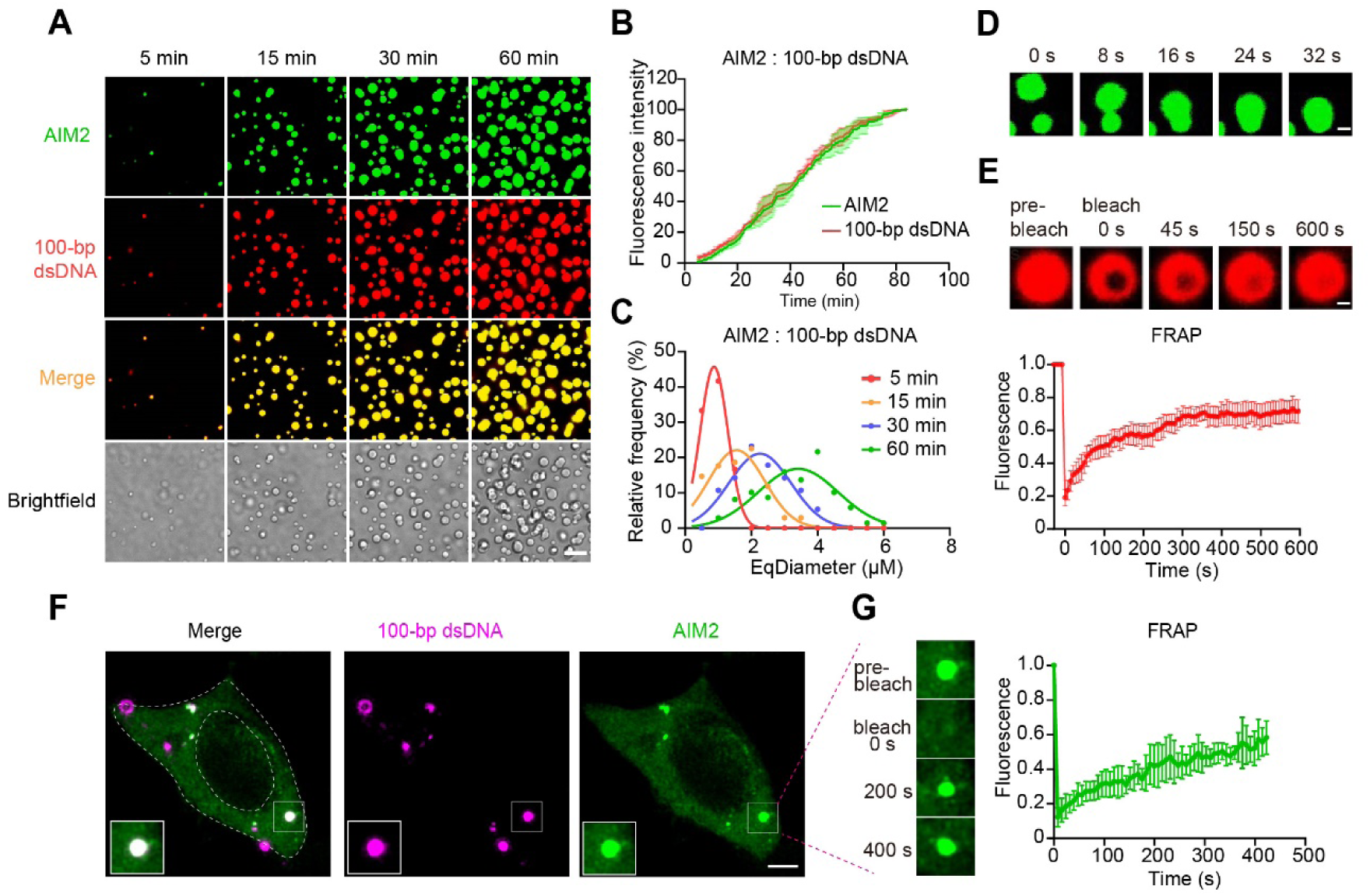
dsDNA induces AIM2 to form liquid-phase condensation *in vitro* and in cells. (A) Time-lapse images of droplets formed by 5 μM AF488-labeled mAIM2 and 1 μM cy3-labeled 100-bp dsDNA in 300 mM NaCl buffer. Scale bar, 10 μm. (B) Fluorescence intensities of mAIM2-DNA droplets formed over the time course of 80 min. The maximal fluorescence intensity was normalized to 100%. Values shown are means ± SD. n=3 images. (C) EqDiameter frequency distribution of mAIM2-DNA droplets that formed as described in (A) at the indicated time points. (D) Time-lapse micrographs of merging mAIM2-DNA droplets that formed as described in (A). Scale bar, 1 μm. (E) FRAP analysis of mAIM2-DNA droplets that formed as described in (A). Top panel: time-lapse images of mAIM2-DNA droplets before and after photobleaching. scale bar, 1 μm. Bottom panel: quantification of FRAP data. Values shown are means ± SD. n=3 condensates. (F) Representative live-cell images of mAIM2-DNA puncta in HEK293T cells expressing GFP-mAIM2 (Tet-on) at 5h after transfecting with 1 μg/mL cy5-labeled 100-bp dsDNA and treating 25 µg/mL doxycycline to induce the expression of GFP-mAIM2. The mAIM2-DNA puncta highlighted by the white box is magnified at the bottom left. scale bar, 4 μm; 4 μm for magnified images. (G) FRAP analysis of mAIM2-DNA puncta that shown in (F). Left panel: time-lapse images of mAIM2-DNA puncta before and after photobleaching. 4 μm for magnified images. Right panel: quantification of FRAP data. Values shown are means ± SD. n=3 puncta.

To better examine AIM2 condensate formation in cells, we generated HEK293T cell lines stably expressing GFP-tagged mAIM2 or hAIM2. Following transfection with cy5-labeled 100-bp dsDNA, both mAIM2 (Figure 1F) and hAIM2 (Figure S1G) readily form puncta with transfected dsDNA in the cytoplasm. FRAP experiments demonstrate robust fluorescence recovery of both mAIM2-DNA and hAIM2-DNA puncta (Figures 1G and S1H), highlighting the dynamic and liquid-like features of AIM2-DNA condensates in cells. Taken together, these results show that DNA induces AIM2 to undergo liquid-liquid phase separation, leading to condensate formation both *in vitro* and in cells, a process conserved between human and mouse proteins.

We further investigated the conditions influencing AIM2 condensate formation. The phase diagram (Figure S1I) shows that AIM2 forms condensates with dsDNA when the concentrations of AIM2 and dsDNA exceeds certain thresholds, with higher concentrations leading to more abundant and larger liquid droplets. Notably, AIM2 does not form robust condensates in the absence of dsDNA, even at relatively high protein concentrations, underscoring the essential role of DNA binding in this process (Figure S1I). Increasing salt concentrations significantly weakens AIM2-DNA phase separation (Figure S1D), indicating that ionic interactions between AIM2 and dsDNA are critical for condensate formation. In addition, the formation of AIM2–DNA droplets is largely inhibited by adding 5% 1,6-hexanediol (Figure S1J). Furthermore, long dsDNA, which provides more binding sites, induces stronger AIM2 phase separation than short dsDNA (Figure S1K), consistent with AIM2 activation requires a certain DNA length (Figure S1L) and also suggesting the critical role of multivalent interactions in this process.

### Multivalent interactions drive AIM2-DNA liquid-phase condensation

To elucidate the molecular mechanisms of AIM2-DNA condensates formation, we performed *in vitro* phase separation assays using wild-type (WT) AIM2 and various truncation and mutation variants (Figure 2A, Table S1 and S2). Compared to WT protein, deletion of PYD (ΔPYD) in mAIM2 nearly abolishes its DNA-induced phase separation ability (Figure 2B). Interestingly, because hAIM2 has a stronger condensation propensity than mAIM2 (Figure 2B and S1F), PYD deletion does not completely eliminate its phase separation but shift it from a gel-like to a typical liquid-like state (Figure 2B). We hypothesized that the essential role of PYD in AIM2-DNA phase separation could be attributed to homotypic PYD-PYD interactions, and indeed, mutating two key residues at the PYD-PYD interface (D15R and Y74R, referred to as PYD_M) produces a similar phenotype to the ΔPYD variant (Figure 2B). Additionally, deleting the HIN domain (ΔHIN) or the entire IDR-HIN region (ΔIDR-HIN) completely abolishes DNA-induced phase separation in both mAIM2 and hAIM2, highlighting the critical role of the HIN domain (Figure 2B). Furthermore, in hAIM2, deletion of the whole PYD-IDR region leads to a more pronounced reduction in phase separation than PYD deletion alone, emphasizing the IDRs essential contribution. Collectively, these results demonstrate that all three major regions of AIM2—PYD, HIN, and IDR—are crucial for providing the multivalent interactions required for AIM2-DNA condensates formation.

**Figure 2.**
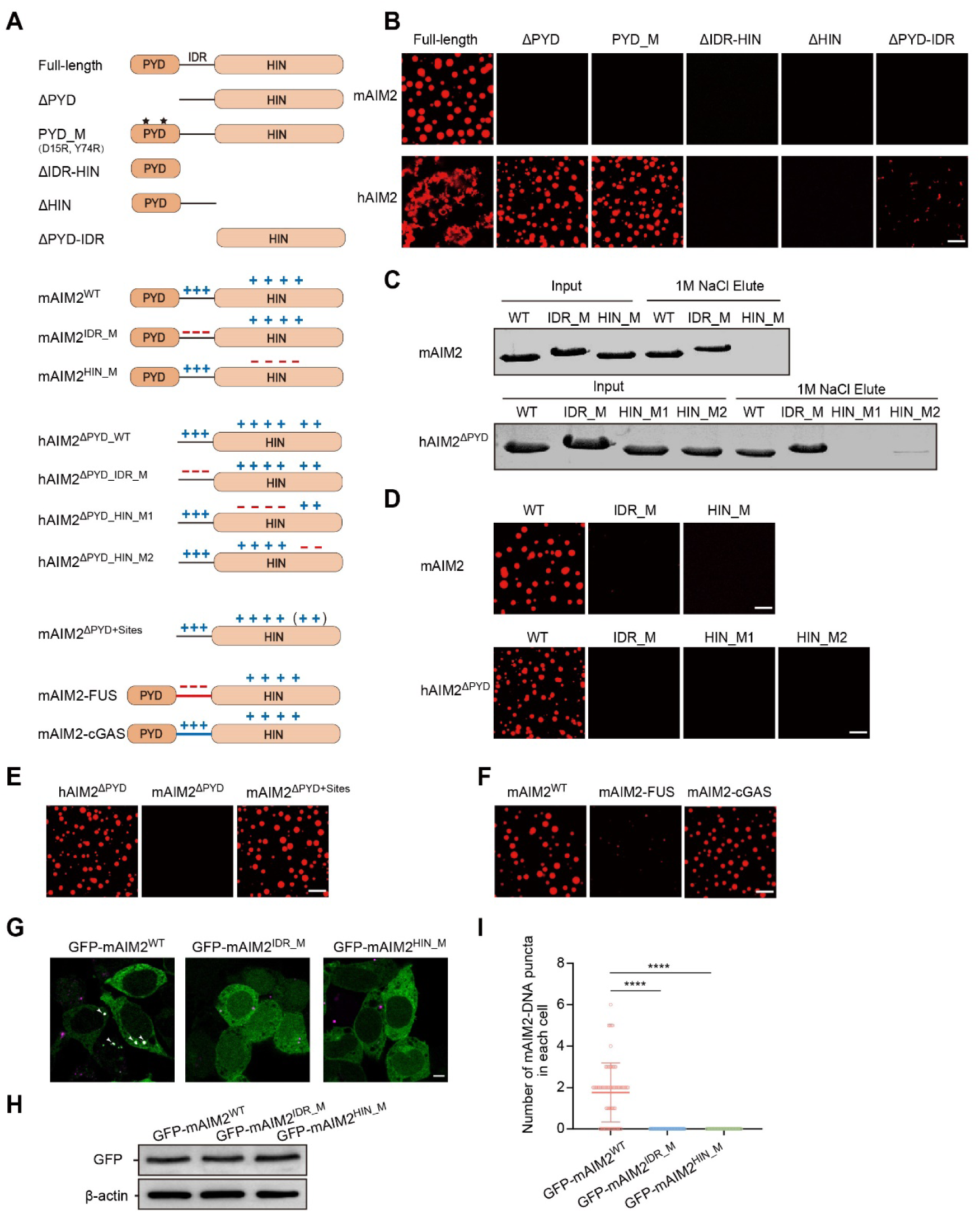
Multivalent interactions drive AIM2-DNA liquid-phase condensation. (A) Schematic of the designed AIM2 constructs. (B) Phase separation assay of indicated AIM2 constructs (5 μM) with cy3-labled 100-bp dsDNA (1 μM) in 150 mM NaCl buffer. Scale bar, 10 μm. (C) dsDNA cellulose beads pull-down assay of mAIM2^WT^ and its mutants (top panel), hAIM2^ΔPYD_WT^ and its mutants (bottom panel). (D) Phase separation assay of mAIM2^WT^ and its mutants (5 μM) with cy3-labled 100-bp dsDNA (1 μM) in 300 mM NaCl buffer (top panel). Phase separation assay of hAIM2^ΔPYD_WT^ and its mutants (5 μM) with cy3-labled 100-bp dsDNA (1 μM) in 150 mM NaCl buffer (bottom panel). Scale bar, 10 μm. (E) Phase separation assay of 5 μM hAIM2^ΔPYD_WT^, mAIM2^ΔPYD_WT^ and mAIM2^ΔPYD+Sites^ with 1 μM cy3-labled 100-bp dsDNA in 150 mM NaCl buffer. Scale bar, 10 μm. (F) Phase separation assay of 5 μM mAIM2^WT^, mAIM2-FUS and mAIM2-cGAS with 1 μM cy3-labled 100-bp dsDNA in 300 mM NaCl buffer. Scale bar, 10 μm. (G) Representative images of mAIM2-DNA puncta (white) in HEK293T cells expressing GFP-mAIM2^WT^, GFP-mAIM2^IDR_M^ or GFP-mAIM2^HIN_M^ (green) at 5h after transfecting with 1 μg/mL cy5-labeled 100-bp dsDNA (magenta) by using lipofectamine 3000 reagent and treating 25 µg/mL doxycycline to induce the expression of GFP-mAIM2^WT^ or its mutants. Arrowheads indicate AIM2-DNA puncta. Scale bar, 5 μm. (H) Immunoblot analysis of the expression of GFP-mAIM2^WT^ and its mutants in (G). (I) Quantification of the number of mAIM2-DNA puncta in each cell that formed as described in (G). n=70. Data are mean ± SD. Statistical analyses were performed by using two-tailed Student’s t test. ****p < 0.0001.

To further understand how the HIN domain influences AIM2-DNA condensates formation, we generated a series of AIM2 mutants targeting potential multivalent interactions with dsDNA (Figure 2A, Table S2). Structural analysis of AIM2^HIN^-DNA complex crystal structures, along with amino acid sequence examination, identifies several positively charged surface residues likely involved in DNA binding (Figures S2A-S2C). Both mAIM2 and hAIM2 share a primary DNA binding site in a similar position, with slight differences in binding orientation (Figures S2A-S2C), which has been considered as the assembly interface in the proposed AIM2 activation model.^32,36^ Interestingly, detailed analysis of the crystal packing lattice reveals two additional DNA binding sites in hAIM2 that are absent in mAIM2 (Figures S2A-S2C). Charge-reversal substitutions at either the primary DNA binding sites (mAIM2^HIN_M^ and hAIM2^ΔPYD_HIN_M1^) or the additional DNA binding sites (hAIM2^ΔPYD_HIN_M2^) dramatically reduce DNA binding (Figures 2C and S2D) and abolish phase condensation (Figure 2D). We hypothesized that the stronger phase separation capacity of hAIM2 compared to mAIM2 may arise from these additional DNA binding sites. Indeed, introducing additional DNA binding sites into the PYD-deleted mAIM2 (mAIM2^ΔPYD+Sites^) restores a phase separation capacity comparable to that of its human counterpart (Figure 2E). These findings show that both the primary and additional DNA binding sites are critical for DNA-induced phase separation and that the additional sites in hAIM2 enhance its condensation propensity relative to mAIM2.

Compared to the proposed AIM2 activation model suggesting that the IDR merely provides flexibility and length between PYD and HIN domains, our results reveal a previously unrecognized role of IDR in AIM2-DNA condensation (Figure 2B). Detailed analysis reveals that both mAIM2 and hAIM2 IDRs harbor multiple positively charged residues (Figure S2A), potentially promoting charge interactions critical for liquid-phase condensation. Charge-reversal mutations at these residues (mAIM2^IDR_M^ and hAIM2^ΔPYD_IDR_M^) practically abolish DNA-induced phase separation (Figure 2D) while minimally affecting DNA binding (Figures 2C and S2D). To further probe the role of IDR, we generated two chimeric proteins, mAIM2-cGAS and mAIM2-FUS, by replacing the mAIM2 IDR with an equally long, positively charged IDR from cGAS or negatively charged IDR from FUS, respectively (Figure 2A). The AIM2-cGAS chimera retains phase separation similar to WT mAIM2, whereas the AIM2-FUS chimera shows a dramatic reduction (Figure 2F). These data highlight the critical role of positively charged residues within the IDR in driving AIM2-DNA liquid-phase condensation.

To validate the importance of these sites in cells, we generated HEK293T cell lines expressing GFP-tagged WT mAIM2 and two condensation-deficient mutants targeting the IDR (mAIM2^IDR_M^) and HIN (mAIM2^HIN_M^) domains. Following transfection with cy5-labeled 100-bp dsDNA, cells expressing mutant AIM2 show no AIM2-DNA puncta formation, in contrast to the robust puncta observed in cells expressing WT protein (Figures 2G-2I). Taken together, both *in vitro* and cellular experiments demonstrate that reducing multivalent protein-DNA or protein-protein interactions weakens AIM2-DNA phase separation, whereas increasing interaction sites enhances its condensation. Moreover, these findings also highlight that, beyond the established PYD-PYD and primary HIN-DNA interfaces, previously underappreciated regions—such as the IDR and additional DNA binding sites within the HIN domain—are also critical for AIM2-DNA liquid-phase condensation.

### AIM2-DNA condensation facilitates inflammasome assembly

Since dsDNA-activated AIM2 recruits ASC and caspase-1 to form inflammasome, we hypothesized that DNA-induced AIM2 condensation may contribute to this assembly process. To test this, we incubated the purified ASC and an enzymatically inactive caspase-1^C284A^ with AIM2-DNA condensates. The results show that ASC is efficiently enriched within the condensates and promotes their solidification (Figures 3A and S3A), while caspase-1 initially localizes to the periphery and then gradually infiltrates the condensates over time (Figures 3A and S3B). In contrast, neither ASC nor caspase-1 is effectively concentrated when incubated with dsDNA and the condensation-deficient mutants (AIM2^IDR_M^ or AIM2^HIN_M^) (Figure 3A). AIM2 phase condensation significantly increases its local concentration, likely promoting homotypic PYD-PYD interactions and the formation of ordered nucleation assemblies. The robust enrichment of ASC and caspase-1 in AIM2-DNA condensates further suggests a potential role in facilitating ASC filament formation and inflammasome assembly. Notably, compared with mAIM2, hAIM2 undergoes phase separation more rapidly, resulting in faster ASC recruitment and nucleation, as supported by the FRET-based kinetic analysis (Figures S3C and S3D).

**Figure 3.**
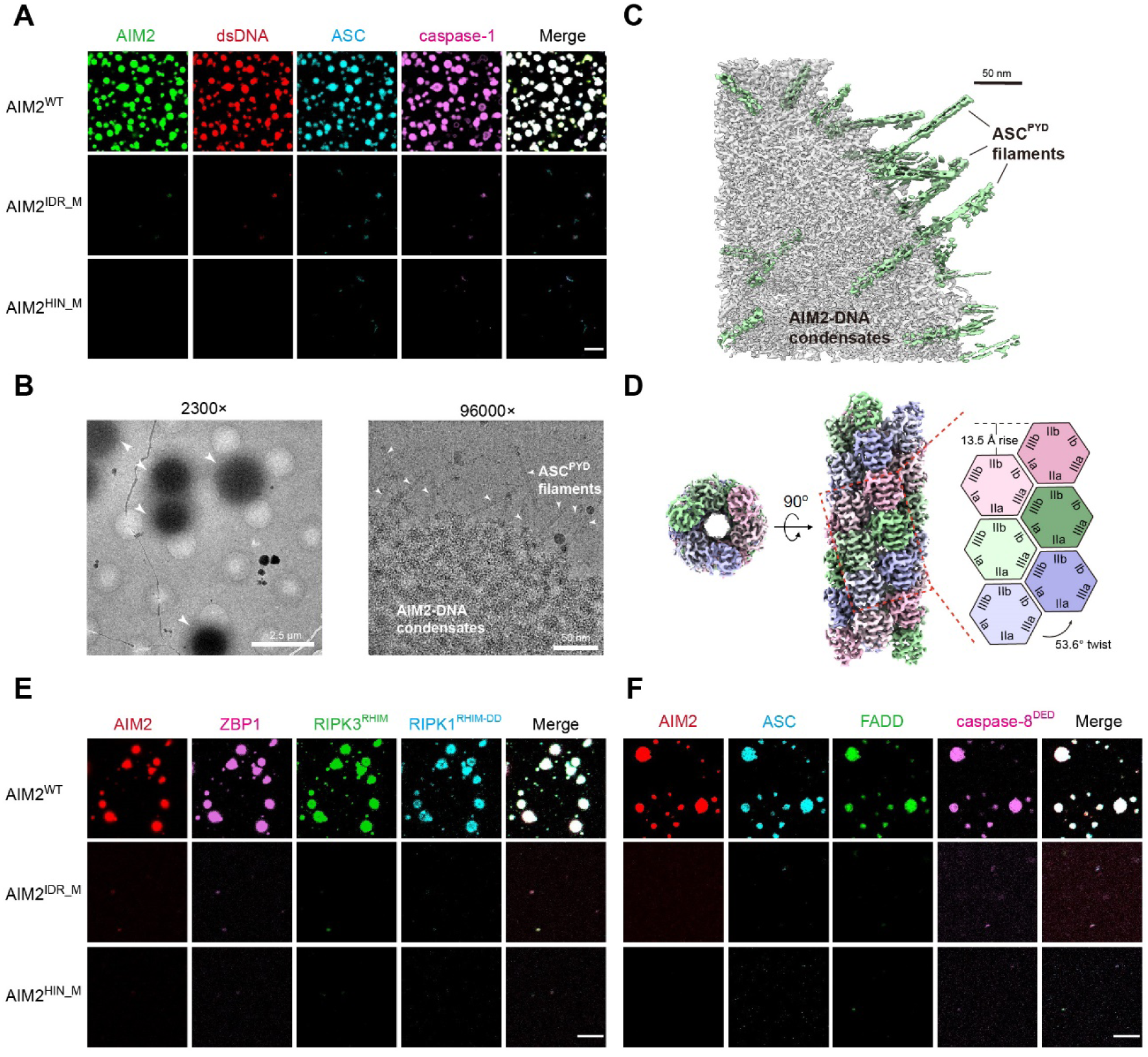
AIM2-DNA condensation facilitates inflammasome and PANoptosome assembly. (A) AIM2-DNA condensates enrich ASC and caspase-1. Incubation 2 μM DyLight405-labeled ASC and 2 μM AF647-labeled caspase-1(C284A) with condensates formed by 5 μM AIM2^WT^ or its mutants and 1 μM cy3-labled 100-bp dsDNA in 150 mM NaCl buffer. Scale bar, 10 μm. (B) Cryo-EM images of AIM2-DNA-ASC^PYD^ sample at indicated magnification. Left panel: 2300×, arrowheads indicate AIM2-DNA condensates. Scale bar, 2.5 μm. Right panel: 96000×, arrowheads indicate the ASC^PYD^ filaments. Scale bar, 50 nm. (C) Cryo-ET of AIM2-DNA-ASC^PYD^ sample. Reconstructed tomogram with 3D rendering is shown, gray volume represents AIM2-DNA condensates, green volume represents ASC^PYD^ filaments. Scale bar, 50 nm. (D) Cryo-EM density map of the ASC^PYD^ filament from AIM2-DNA-ASC^PYD^ sample. Left panel: Top view. Middel panel: side view. Right panel: 2D schematic diagram of the ASC^PYD^ assembly. Three types of asymmetric interaction interfaces and helical indexing are shown. (E-F) AIM2-DNA condensates enrich multiple PANoptosome components. (E) Co-incubate 2 μM AF647-labeled ZBP1, 2 μM GFP-RIPK3^RHIM^, 2 μM DyLight 405-labeled RIPK1^RHIM-DD^ with 5 μM AF555-labeled AIM2 and 1 μM 100-bp GC-DNA in 150 mM NaCl buffer. (F) Co-incubate 2 μM DyLight 405-labeled ASC, 2 μM AF488-labeled FADD, 2 μM AF647-labeled caspase-8^DED^ with 5 μM AF555-labeled AIM2 and 1 μM 100-bp GC-DNA in 150 mM NaCl buffer. Scale bar, 5 μm.

Given that ASC^PYD^ filament formation represents a hallmark and critical step of AIM2-mediated inflammasome activation, we intended to verify whether AIM2-DNA condensates promote this process. We generated cryo-electron tomography (cryo-ET) samples by incubating ASC^PYD^ with AIM2-DNA condensates and collected data by tilting the sample stage from –50° to +50°. The reconstructed tomogram shows that numerous ASC^PYD^ filaments extending from AIM2-DNA condensates (Figures 3B and 3C, Video S1). Furthermore, to obtain high-resolution structural information, we performed cryo-electron microscopy (cryo-EM) analysis for these samples, obtaining a 2.7 Å reconstruction of the ASC^PYD^ filament (Figures 3D and S3E-S3I, Table S3). The atomic model reveals a right-handed helical structure with a twist angle of 53.6° and an axial rise of 13.5 Å, mediated by three major interaction interfaces of each ASC^PYD^ subunit (Figure 3D), consistent with previously reported activated ASC^PYD^ filament structures.^5^ Together, these data show that DNA-induced AIM2 phase separation promotes inflammasome assembly by efficiently enriching and nucleating ASC and caspase-1.

### AIM2-DNA condensation facilitates PANoptosome assembly

In addition to mediating inflammasome assembly, AIM2 has also been shown to form PANoptosome with other proteins, including ZBP1, RIPK1, RIPK3, FADD, and caspase-8, to mediate inflammatory PANoptosis combining pyroptosis, apoptosis, and necroptosis.^15–18^ To assess whether DNA-induced AIM2 phase separation facilitates PANoptosome assembly, we expressed and purified ASC, ZBP1, RIPK1^RHIM-DD^, RIPK3^RHIM^, FADD, and caspase-8^DED^, and simultaneously incubated them with dsDNA and either WT AIM2 or its condensation-deficient mutants. The results show that all these proteins are efficiently enriched into AIM2-DNA condensates (Figures 3E and 3F), consistent with these protein colocalized with AIM2 puncta in cells as previous study reported.^15^ However, no effective enrichment was observed when these proteins were incubated with dsDNA and AIM2^IDR_M^ or AIM2^HIN_M^ mutants (Figures 3E and 3F). These molecular evidences align well with prior cellular colocalization experiments and mechanistically explain why AIM2 deficiency disrupts PANoptosis activation.

Overall, these findings demonstrate that AIM2-DNA condensation not only promotes inflammasome assembly but also contributes to PANoptosome assembly by recruiting and enriching associated factors.

### AIM2-DNA condensation promotes broad immune activation

Since AIM2-DNA phase separation drives the assembly of both inflammasome and PANoptosome, we reasoned that this condensation process may also play a critical role in downstream immune activation across multiple cell death pathways. To verify this hypothesis, we generated transgenic mice harboring either *Aim2^WT^* or the condensation-deficient mutants (*Aim2^IDR_M^* or *Aim2^HIN_M^*) using the CRISPR-Cas9 method (Figures S4A and S4B). Bone marrow-derived macrophages (BMDMs) isolated from these mice were used for inflammasome and PANoptosome activation assays (Figure 4A). Upon infection with *Francisella novicida* (*F. novicida*), a Gram-negative bacterium known to replicate in the cytoplasm and activate AIM2,^19–21^ *Aim2^WT^*BMDMs exhibit significantly more AIM2-ASC coenriched puncta compared to *Aim2^IDR_M^*or *Aim2^HIN_M^* BMDMs (Figures 4B and 4C). Correspondingly, *Aim2^WT^*BMDMs show robust caspase-1 and GSDMD cleavage and high IL-18 release, whereas these activation markers are markedly reduced in the mutant BMDMs (Figures 4D and 4E). In line with *F. novicida* infection, similar results were observed following poly(dA:dT) stimulation (Figures 4F and 4G). In addition, we assessed apoptosis and necroptosis activation by evaluating caspase-8 and caspase-3 cleavage, as well as RIPK3 and MLKL phosphorylation, in BMDMs infected with *F. novicida*. Compared to *Aim2^WT^* BMDMs, the activation of these apoptosis- and necroptosis-associated proteins is significantly diminished in *Aim2^IDR_M^* and *Aim2^HIN_M^*BMDMs (Figures 4H and 4I). Consistent with these activation markers, the overall cell death induced by *F. novicida* infection in *Aim2^WT^* BMDMs is dramatically higher than in the mutant groups (Figures 4J and 4K). Taken together, these findings demonstrate that AIM2-DNA condensation is not only critical for the efficient assembly of inflammasome and PANoptosome but also plays an essential role in immune activation across pyroptotic, apoptotic, and necroptotic pathways.

**Figure 4.**
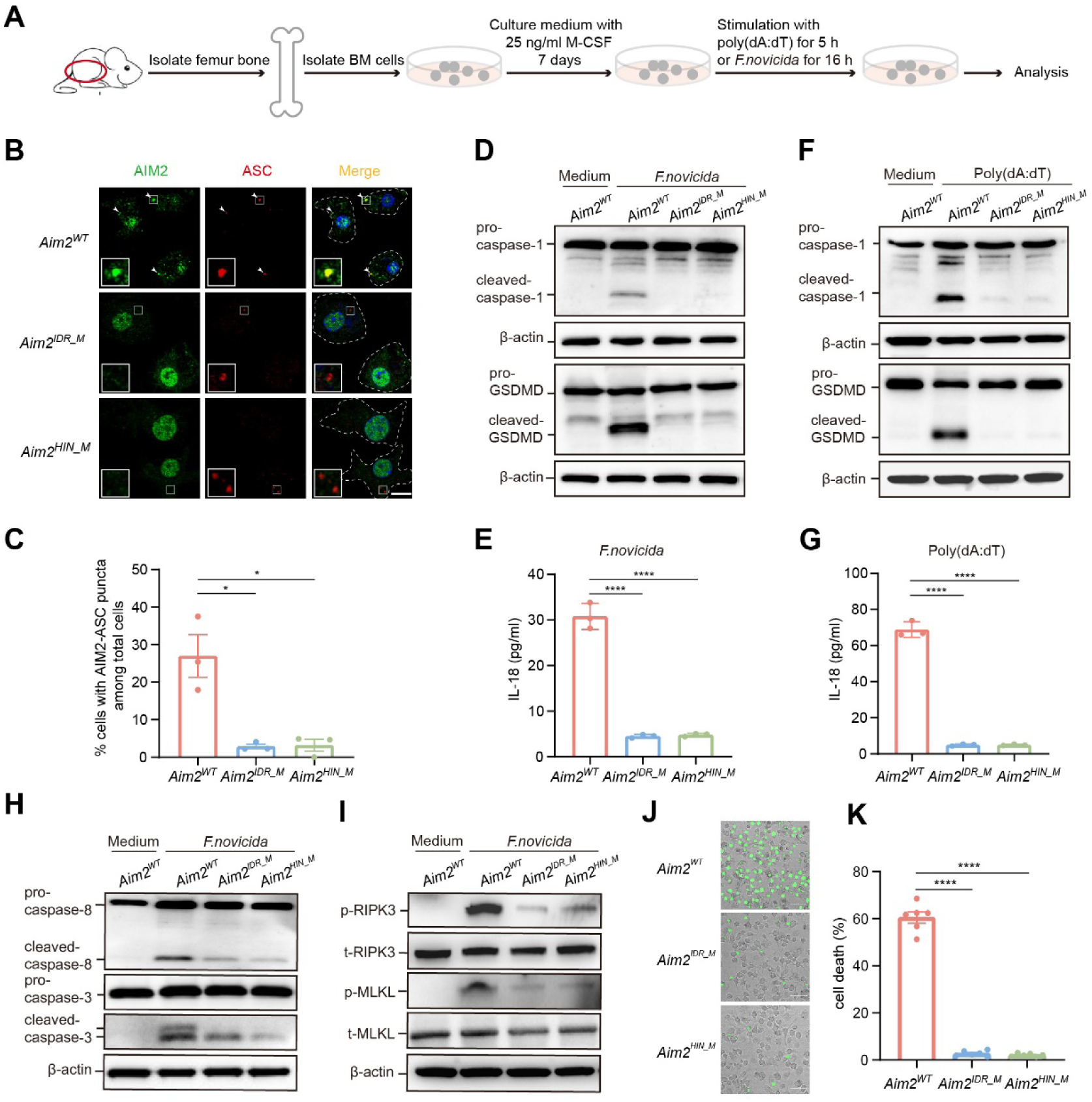
AIM2-DNA condensation promotes broad immune activation. (A) Schematic diagram of BMDMs were isolated from mice and cultured to perform assay. (B) Immunofluorescence images of *Aim2^WT^*, *Aim2^IDR_M^* and *Aim2^HIN_M^* BMDMs at 16 h after *F. novicida* infection. Arrowheads indicate the AIM2-ASC puncta. Scale bar, 10 μm. 4 μm for magnified images. (C) Quantification of the percentage of cells with AIM2-ASC puncta among all cells from (B). These data are representative of three independent experiments. Data are mean ± SEM. *Aim2^WT^*, n=134; *Aim2^IDR_M^*, n=130; *Aim2^HIN_M^*, n=136. Statistical analyses were performed by using two-tailed Student’s t test. *p < 0.05. (D) Immunoblot analysis of pro- and cleaved-caspase-1, pro- and cleaved-GSDMD in *Aim2^WT^*, *Aim2^IDR_M^* and *Aim2^HIN_M^* BMDMs at 16 h after *F. novicida* infection. (E) Cell supernatant IL-18 levels in *Aim2^WT^*, *Aim2^IDR_M^* and *Aim2^HIN_M^* BMDMs at 16 h after *F. novicida* infection. These data from three independent experiments. Data are mean ± SEM. Statistical analyses were performed by using two-tailed Student’s t test. ****p < 0.0001. (F) Immunoblot analysis of pro- and cleaved-caspase-1, pro- and cleaved-GSDMD in *Aim2^WT^*, *Aim2^IDR_M^* and *Aim2^HIN_M^* BMDMs at 5 h after stimulation with poly(dA:dT). (G) Cell supernatant IL-18 levels in *Aim2^WT^*, *Aim2^IDR_M^* and *Aim2^HIN_M^* BMDMs at 5 h after stimulation with poly(dA:dT). These data from three independent experiments. Data are mean ± SEM. Statistical analyses were performed by using two-tailed Student’s t test. ****p < 0.0001. (H) Immunoblot analysis of pro- and cleaved-caspase-8, pro- and cleaved-caspase-3 in *Aim2^WT^*, *Aim2^IDR_M^*and *Aim2^HIN_M^* BMDMs at 16 h after *F. novicida* infection. (I) Immunoblot analysis of phosphorylated RIPK3 (p-RIPK3), total RIPK3 (t-RIPK3), phosphorylated MLKL (p-MLKL), total MLKL (t-MLKL) in *Aim2^WT^*, *Aim2^IDR_M^* and *Aim2^HIN_M^*BMDMs at 16 h after *F. novicida* infection. (J) Cell death in *Aim2^WT^*, *Aim2^IDR_M^*and *Aim2^HIN_M^* BMDMs at 16 h after *F. novicida* infection. Cell death was measured by SYTOX Green uptake assay. Green indicates dead cells. Scale bar, 50 μm. (K) Quantification of cell death from (J). These data from three independent experiments. Data are mean ± SEM. Statistical analyses were performed by using two-tailed Student’s t test. ****p < 0.0001.

### AIM2-DNA condensation promotes *in vivo* anti-infection defense

Given the essential role of AIM2-DNA condensation in activating multiple cell death pathways (Figure 4) and the established link between AIM2 and infection defense,^15,19,20^ we hypothesized that this condensation process is crucial for *in vivo* protection against pathogen infection. To test this, we infected *Aim2^WT^* mice and condensation-deficient mutant mice (*Aim2^IDR_M^* and *Aim2^HIN_M^*) with *F. novicida* and monitored body weight changes, survival, and bacterial loads (Figure 5A). Infection with *F. novicida* results in significantly higher mortality in *Aim2^IDR_M^*and *Aim2^HIN_M^* mice compared to *Aim2^WT^* mice (Figure 5B). All *Aim2^HIN_M^* mice succumbed within 6 days, and all *Aim2^IDR_M^*mice died within 9 days, whereas 80% of *Aim2^WT^* mice survived beyond 14 days post-infection (Figure 5B). Correspondingly, mutant mice exhibited more severe weight loss than *Aim2^WT^* mice (Figure S4C). While *Aim2^WT^* mice regained body weight after an initial decline, mutant mice failed to recover (Figure S4C). Consistently, bacterial loads in the lung, spleen, and liver of *Aim2^IDR_M^* and *Aim2^HIN_M^*mice are significantly higher than those in *Aim2^WT^* mice at 48 hours post-infection (Figure 5C). Together, these data demonstrate that the loss of AIM2-DNA condensation compromises host defense, highlighting its indispensable role in mounting an effective immune response against infection.

**Figure 5.**
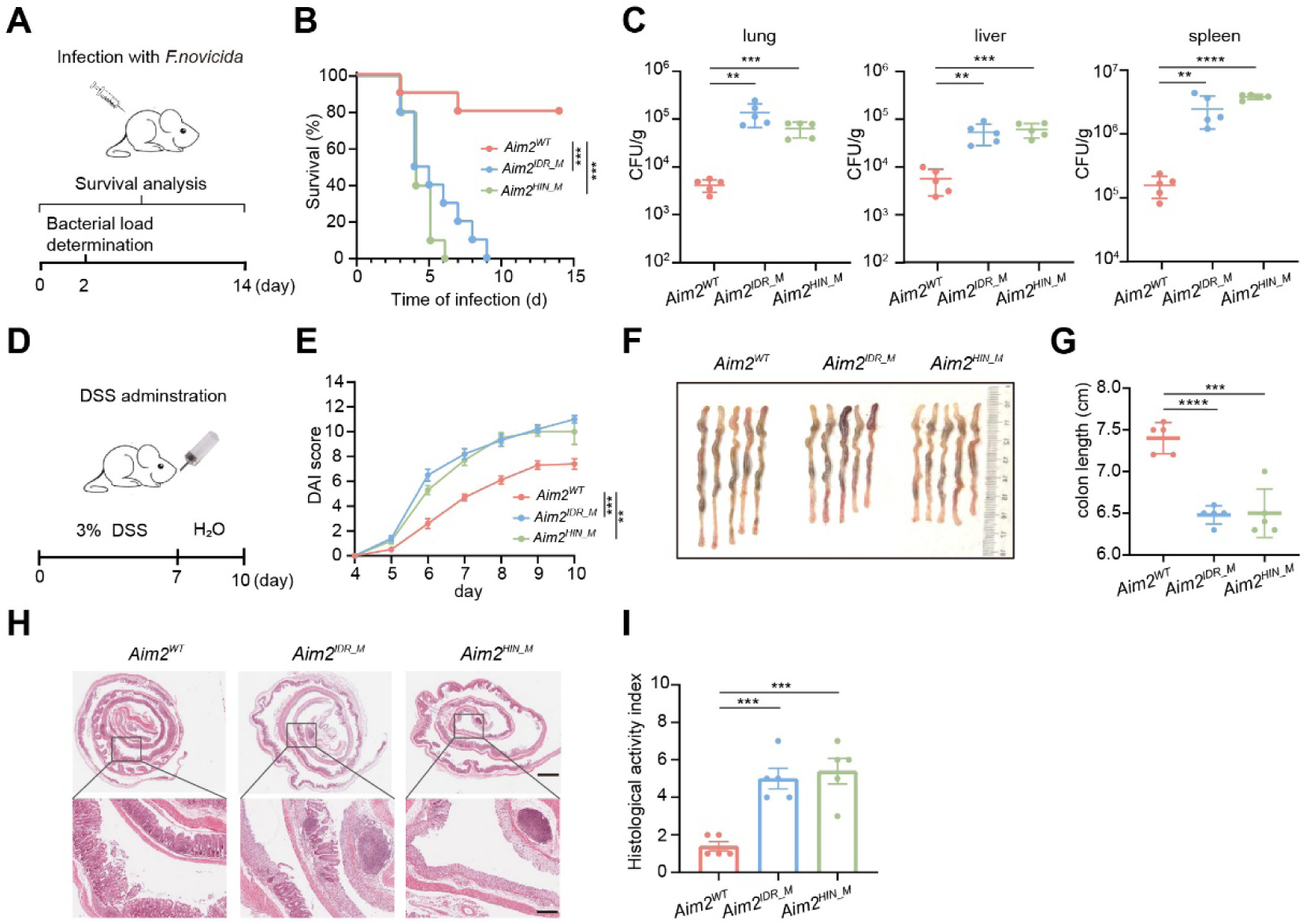
AIM2-DNA condensation promotes *in vivo* anti-infection defense and maintains intestinal homeostasis. (A) Schematic diagram of mice infection assay. (B) Survival of *Aim2^WT^*, *Aim2^IDR_M^*and *Aim2^HIN_M^* mice (10 mice per group) infected with 5×10^5^ CFU of *F. novicida* per mouse. Statistical analyses were performed by using log-rank (Mantel-Cox) test. ***p < 0.001. (C) Bacterial load in lung, liver, spleen obtained from *Aim2^WT^*, *Aim2^IDR_M^* and *Aim2^HIN_M^* mice at 48 h after infection with 5×10^5^ CFU of *F. novicida* per mouse. Data are mean ± SEM, n=5 mice per group. Statistical analyses were performed by using a two-tailed Student’s t test. ****p < 0.0001, ***p < 0.001, **p < 0.01. (D) Schematic diagram of DSS administration assay. (E) DAI score was evaluated daily after DSS administration. Data are mean ± SEM, n=5 mice per group. Statistical analyses were performed by using two-way ANOVA. ***p < 0.001, **p < 0.01. (F) Pictures of colon of *Aim2^WT^*, *Aim2^IDR_M^* and *Aim2^HIN_M^* mice at day 10 after DSS administration. (G) The length of colon in (F). Data are mean ± SEM, n=5 mice per group. Statistical analyses were performed by using a two-tailed Student’s t test. ****p < 0.0001, ***p < 0.001. (H) Representative H&E images of Swiss roll intestine. scale bar, 1.5 mm. The representative region highlighted by the black box is magnified. scale bar, 250 μm. (I) Histological activity index in (H) was evaluated. Data are mean ± SEM, n=5 mice per group. Statistical analyses were performed by using two-tailed Student’s t test. ***p < 0.001.

### AIM2-DNA condensation maintains intestinal homeostasis

AIM2 has also been implicated in maintaining intestinal homeostasis by mediating epithelial antimicrobial host defense.^42,43^ To investigate whether AIM2-DNA phase separation contributes to this process, we established a DSS (dextran sodium sulfate)-induced acute colitis model and evaluated its effects in WT and mutant mice (Figure 5D). Compared to *Aim2^WT^* mice, *Aim2^IDR_M^* and *Aim2^HIN_M^*mice exhibit markedly higher Disease Activity Index (DAI) scores following DSS administration (Figure 5E). In addition, *Aim2^IDR_M^* and *Aim2^HIN_M^*mice show more severe colonic shortening than *Aim2^WT^* mice at day 10 post-DSS treatment (Figures 5F and 5G). Histological assessment further reveals extensive crypt and goblet cell loss, along with increased inflammation infiltration in *Aim2^IDR_M^* and *Aim2^HIN_M^* mice compared to *Aim2^WT^* mice (Figures 5H and 5I). These results indicate that AIM2 condensation deficiency aggravates colitis severity upon DSS administration, underscoring the critical role of DNA-induced AIM2 phase separation in maintaining intestinal homeostasis.

### AIM2-DNA condensation is modulated by host and pathogen factors

Given the critical role of DNA-induced AIM2 condensation in immune activation and various *in vivo* functions, this process is likely a key regulatory target for different host and pathogen factors. p202, a mouse-encoded AIM2-like protein containing two HIN domains but lacking a PYD, has been reported to interact with AIM2 and suppress its activation.^4,32,33^ To determine whether p202 influences AIM2-DNA phase separation, we incubated purified p202 with AIM2 and dsDNA. Compared to the buffer control, p202 significantly disrupts AIM2-DNA condensates, instead forming a few residual irregular aggregates with AIM2 and dsDNA (Figure 6A). Consistently, in HEK293T cells co-expressing AIM2, ASC, and p202, the formation of AIM2-ASC coenriched puncta is markedly reduced compared to cells lacking p202 expression (Figures 6B, 6C and S5A). Although p202 co-localizes with AIM2—mirroring the *in vitro* observations— ASC is not effectively recruited to AIM2 condensates (Figure 6B). Furthermore, replacing WT p202 with a DNA-binding-deficient mutant (p202^M^) restores the formation of AIM2-ASC coenriched puncta (Figures 6B, 6C and S5A). These findings indicate that p202 disrupts AIM2-DNA phase separation in a DNA-binding-dependent manner, thereby suppressing AIM2 activation. However, other cytosolic DNA-binding proteins such as cGAS and ZCCHC3 do not perturb AIM2 condensation (Figures S5B and S5C), suggesting that p202-mediated disruption likely requires not only DNA binding but also a direct interaction with AIM2.

**Figure 6.**
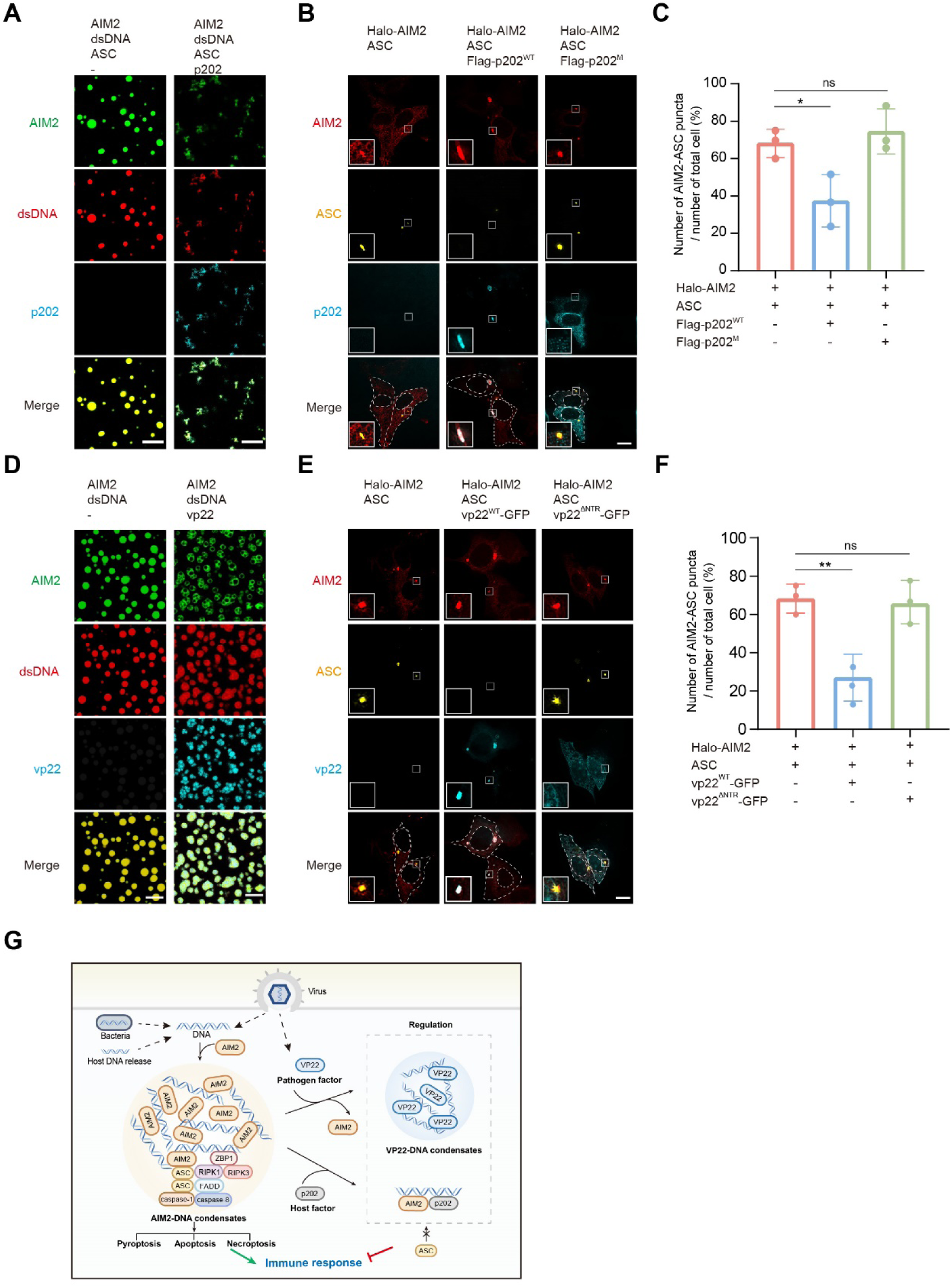
Host- and pathogen-mediated regulation of AIM2-DNA phase separation. (A) Images of 5 μM AF488-labeled AIM2, 1 μM cy3-labeled 100-bp dsDNA incubated with buffer (left panel) or 10 μM Dylight 405-labeled p202 (right panel) in 150 mM NaCl buffer. Scale bar, 10 μm. (B) Immunofluorescence images of HEK293T cells transfected with indicated plasmids for 12 h. Scale bar, 10 μm. 5 μm for magnified images. (C) Quantification of AIM2-ASC puncta among total cells in (B). Data are mean ± SD. Statistical analyses were performed using a two-tailed Student’s t test. *p < 0.05. NS, not significant. (D) Images of 5 μM AF488-labeled AIM2 and 1 μM cy3-labeled 100-bp dsDNA incubated with buffer (left panel) or 10 μM AF647-labeled VP22 (right panel) in 150 mM NaCl buffer. Scale bar, 10 μm. (E) Immunofluorescence images of HEK293T cells transfected with indicated plasmids for 12 h. Scale bar, 10 μm. 5 μm for magnified images. (F) Quantification of AIM2-ASC puncta among total cells in (E). Data are mean ± SD. Statistical analyses were performed using a two-tailed Student’s t test. **p < 0.01. NS, not significant. (G) Model of AIM2-DNA liquid-phase condensation and its function in immune activation and regulation.

VP22, a tegument protein from HSV-1, has also been reported to inhibit AIM2 inflammasome activation by preventing AIM2 oligomerization.^34,35^ To assess whether VP22 directly interacts with AIM2, we performed *in vitro* pull-down assays using purified proteins, with contaminating nucleic acids removed, but detected no clear interaction under physiological buffer conditions (Figure S5D). We next examined whether VP22 interferes with AIM2-DNA phase separation. Co-incubation of VP22, AIM2, and dsDNA shows that VP22 robustly occupies a large portion of the AIM2-DNA condensates, forming its own phase separation with dsDNA while excluding AIM2 (Figure 6D). Furthermore, when VP22 first forms condensates with DNA prior to AIM2 addition, AIM2 is sequestered to the periphery of VP22-DNA condensates (Figure S5E). In both cases, AIM2-DNA phase separation is notably reduced. To validate these results in cells, we transfected HEK293T cells with plasmids that expressing AIM2, ASC, and VP22. Compared to cells lacking VP22, those expressing VP22 exhibit a significant reduction in AIM2-ASC puncta formation (Figures 6E, 6F and S5G). Although AIM2 and VP22 initially colocalize at 12 hours post-transfection (Figure 6E), they exhibit distinct spatial separation by 24 hours (Figure S5F). Importantly, at both time points, ASC fails to be recruited to AIM2 foci (Figures 6E and S5F), suggesting that VP22 effectively disrupts AIM2-mediated inflammasome assembly. To further delineate the role of VP22 phase separation, we employed VP22^ΔNTR^, a phase separation-defective mutant.^44^ Expression of VP22^ΔNTR^ restores AIM2-ASC puncta formation (Figures 6E, 6F and S5G), indicating that VP22 relies on its phase separation ability to disrupt AIM2-DNA condensates, thereby inhibiting AIM2 activation.

## DISCUSSION

Our study identifies a previously unrecognized mechanism of AIM2-mediated immune activation, wherein aberrant cytosolic dsDNA from infection or cellular stress induces AIM2 liquid-liquid phase separation to form biomolecular condensates (Figure 6G). These condensates act as dynamic multifunctional platforms, assembling the AIM2 inflammasome by recruiting ASC and caspase-1, while also integrating ZBP1, RIPK1, RIPK3, FADD, and caspase-8 to form AIM2 PANoptosome, thereby triggering pyroptosis, apoptosis, and necroptosis. In addition, AIM2 phase separation may serve as a “checkpoint“—enabling rapid immune activation when cytosolic DNA reaches a physiologically meaningful length threshold while preventing inadvertent activation by shorter DNA fragments. Beyond its role in immune activation, AIM2-DNA condensates also serve as key regulatory hubs targeted by host- (e.g., p202) and pathogen-derived (e.g., VP22) factors, balancing immune homeostasis or facilitating immune evasion. *In vivo* experiments further underscore the essential role of AIM2-DNA condensation in mounting effective antimicrobial defenses and maintaining intestinal homeostasis. Notably, condensation-deficient AIM2 mutants fail to assemble immune complexes, initiate cell death pathways, or exert protective functions *in vivo*, highlighting the physiological importance of this phase separation process in innate immunity.

Additionally, our investigation into the molecular basis of AIM2-DNA condensation reveals that this process is driven by multivalent protein-DNA and protein-protein interactions, with the PYD, HIN, and IDR domains each being indispensable for condensate formation. PYD-PYD homotypic interactions not only promote phase separation but also facilitate the hierarchical assembly of PYD-dependent immune complexes, positioning condensates as platforms for signal amplification. The HIN domain, beyond its primary DNA-binding site, contains two additional DNA-binding sites in hAIM2 compared to its murine counterpart, enhancing DNA avidity and phase separation. This species-specific divergence likely optimizes innate immune responses by enabling cooperative DNA engagement and stabilizing higher-order AIM2-DNA assemblies. Meanwhile, the IDR, previously considered merely a flexible linker, emerges as a key modulator of condensate formation. Disrupting its charge distribution impairs phase separation without affecting DNA binding, suggesting it mediates weak multivalent interactions crucial for condensate assembly while preserving DNA recognition. Notably, IDR mutations that selectively disrupt phase separation severely compromise immune activation, infection resistance, and intestinal homeostasis, highlighting the instructive and causal role of phase separation in AIM2 signaling.

The currently identified AIM2-DNA condensation mechanism offers a more comprehensive understanding of AIM2-mediated immune activation compared to previously predicted models. Phase separation allows AIM2 to efficiently concentrate at dsDNA within the vast cytosolic space, addressing the challenge of how relatively low AIM2 levels converge on DNA targets. This process not only ensures DNA recognition but also clarifies the previously unresolved DNA length requirement for AIM2 activation. Furthermore, phase separation significantly increases the local concentration of AIM2 and DNA, enhancing the recruitment of downstream immune factors and promoting inflammasome and PANoptosome assembly. Additionally, previously overlooked sites within the IDR and HIN domains have been identified as critical for both phase separation and AIM2 function, providing new insights into how AIM2 interacts with DNA and assembles immune complexes. Moreover, this model uncovers a novel immune regulatory mechanism, where both host and viral proteins can modulate the condensation process. Notably, the viral protein VP22, despite not having evolved strong direct interactions with AIM2, can still fine-tune immune activation by influencing AIM2-DNA phase separation. This mechanism mirrors the regulatory role of this protein family in modulating cGAS-DNA phase separation.^44,45^

As another key cytosolic DNA sensor, cGAS has also been shown to undergo phase separation with DNA.^45^ This raises the question of how AIM2 and cGAS behave when aberrant DNA accumulates in the same cellular environment. To explore this, we examined their subcellular localization following DNA stimulation in BMDMs and THP1 cells, both of which express endogenous AIM2 and cGAS. We observed that AIM2 and cGAS co-localize with DNA and form condensates (Figure S5H). Consistent with the cellular results, cGAS also co-partitions with AIM2-DNA condensates *in vitro* (Figure S5C). These observations suggest that the two sensors can coexist within shared DNA-rich condensates, although whether they influence each others condensation dynamics or downstream signaling remains an open question for future investigation.

In summary, this study uncovers a novel mechanism of AIM2-DNA condensation in immune activation and regulation, offering new perspectives on the physiological and pathological functions of AIM2, and providing potential therapeutic opportunities for targeting the condensation process to modulate immune responses.

## Data Availability

The atomic coordinate of ASC^PYD^ filament has been deposited at the Protein Data Bank under accession numbers 9U8K. The cryo-EM density map has been deposited at the Electron Microscopy Data Bank under accession numbers EMD-63955.

## ACKNOWLEDGMENTS

Cryo-ET and Cryo-EM data collection was carried out at the Center for Biological Imaging, Core Facilities for Protein Science at the Institute of Biophysics, Chinese Academy of Sciences. The computation work was performed using High-performance computing resources, the Center for Biological Imaging, Institute of Biophysics, Chinese Academy of Science. We thank B. Zhu, S. Li, X. Huang, G. Ji, F. Sun and other staff members at the Center for Biological Imaging for their support in data collection; C. Qi and Y. Feng for analyzing cryo-ET data. We also thank Cryo-Electron Microscopy Platform of Medical Science and Technology Innovation Center of Shandong First Medical University for the support of Cryo-EM data collection. We thank Y. Teng, Q. Bian, X. Jia, and Y. Xu for helping with confocal imaging; J. Jia, S. Meng and X. Wang for helping with FACS; We thank Prof. Jingren Zhang for providing *F. novicida* strain U112. This work was supported by grants from National Science and Technology Major Project of China (2025ZD01904303), National Natural Science Foundation of China (32325028 & 32130057, 32571417), National Key R&D Program of China (2024YFA1307400), Strategic Priority Research Program (XDB1310000), Beijing Natural Science Foundation (Z220018), Basic Research Program Based on Major Scientific Infrastructures-CAS (JZHKYPT-2021-05), CAS Project for Young Scientists in Basic Research (YSBR-074).

## CONTRIBUTIONS

Q.L., X.G., H.Y., and Z.L performed experiments with the help of Z.Z., C.Z., K.S., Y.G., M.S., S.L. and Q.J.; P.B., D.L., and P.G. designed and directed the research; Q.L. and P.G. wrote the manuscript with the help of all authors.

## Methods

### Mice

All mice were bred and housed under specific pathogen-Free facilities at the Institute of Biophysics, Chinese Academy of Sciences. Mouse maintenance and procedures were conducted in accordance with the ethical guidelines for animal research and were approved by the Biomedical Research Ethics Committee of the Institute of Biophysics, Chinese Academy of Science. All mice were maintained with 5 mice per cage in a specific-pathogen free facility with a 12h:12h light:dark cycle at 21-23°C. The following mouse strains were used in this study: *Aim2^WT^*, *Aim2^IDR_M^* and *Aim2^HIN_M^* mice on the C57BL/6 background were obtained from GemPharmatech corporation.

### Mammalian cell lines

BMDMs and HEK293T were cultured in DMEM media (GBICO) supplemented with 10% fetal bovine serum (Thermo Fisher Scientific) at 37°C and 5% CO2. For BMDMs isolation and culture, primary murine bone marrow cells were flushed from tibias and femurs with chilled DMEM and cultured for 7 days in DMEM (C11885500BT, Gibco) supplemented with 10% FBS and 25 ng/ml M-CSF (MCE) to generate BMDMs. For doxycycline-inducible HEK293T stable cell lines, 10% Tet-System Approved FBS (Biological Industries) was added to the DMEM media instead of fetal bovine serum. THP-1 cells were cultured in RPMI media (GBICO) supplemented with 10% fetal bovine serum (Thermo Fisher Scientific) at 37°C and 5% CO2. Before stimulation, THP-1 cells were differentiated for 24 h with 1 μM PMA. All media was supplemented with 1% penicillin-streptomycin.

### Escherichia coli strains

*E.coli* BL21(DE3) cells (Tsingke) were used to express ASC, ASC^PYD^, caspase-1, FADD, caspase-8^DED^, RIPK1^RHIM-DD^, RIPK3^RHIM^-GFP, ZBP1, VP22, p202, AIM2 and its mutants.

### Protein Expression and Purification

*Aim2* and its constructs, *Asc*, *Caspase-1*, *Fadd*, *Caspase-8^DED^*, *Ripk1^RHIM-DD^* and *Ripk3^RHIM^-gfp* were inserted into pET28a plasmid with N-terminal TEV protease cleavable His6-MBP tag. *p202* was inserted into pET28a plasmid with C-terminal TEV protease cleavable His6-MBP tag. *vp22* was inserted into pRSFDUET-1 plasmid with N-terminal His6-SUMO tag. These proteins were expressed in BL21(DE3) cell strain. The cells were grown at 37°C until OD600 reached 0.8. The temperature was then shifted to 20°C and the cells were induced with 0.3 mM IPTG overnight. *Zbp1* was inserted into pFASTBAC plasmid with N-terminal TEV protease cleavable His6-MBP tag and was expressed in SF9 for 3 days. After cells were harvested by centrifugation and resuspended in buffer A (50 mM Tris-HCl pH 7.5, 500mM NaCl, 20 mM imidazole, 5 mM β-mercaptoethanol, 10% glycerol), cells were lysed by French Press and lysates were cleared by centrifugation. The supernatant was loaded onto a Ni-NTA affinity column (GE Healthcare). Sample was eluted with buffer B (50 mM Tris-HCl pH 7.5, 500mM NaCl, 500 mM imidazole, 5 mM β-mercaptoethanol, 10% glycerol). AIM2 and its constructs, p202, VP22 and ZBP1 were further applied to a 5-mL Hitrap Heparin column (GE Healthcare) with a linear NaCl gradient from 150 mM to 1 M. ASC, caspase-1 were further applied to a 5-mL Hitrap Q column (GE Healthcare) with a linear NaCl gradient from 150 mM to 1 M. Then samples were subjected to gel filtration on a Superdex G200 10/300 GL size-exclusion column (GE Healthcare) equilibrated with buffer C (20 mM Hepes pH 7.5, 300 mM NaCl, 1 mM DTT). The peak sample was concentrated, measured and stored at −80°C before use.

### Proteins Labeling, Phase Separation Assay and FRAP assay *in vitro*

Purified proteins were labeled with corresponding dyes by using Alexa Fluor™ Protein Labeling Kit (Thermo Fisher Scientific) according to the manufacturer’s protocols. Phase separation assay performed by mixing indicated proteins (2% dyes labeled), indicated dsDNA (2% cy3 labeled) and 0.2 mg/ml TEV, in a final volume of 50 ml at room temperature. Then mixing samples were loaded into 96-well plates (Corning) pre-coated with 20 mg/mL BSA (Sigma) to be imaged by a Nikon A1R+ confocal microscope with an oil immersion 60× objective lens. Concentration of salt, proteins and dsDNA are indicated in the Fig. legends. For time-lapse imaging, images were acquired with 7 s interval in 60 min. For FRAP assay, the droplets were imaged with 7 s interval for 21 s and then photobleaching was performed with a 561 nm laser. After bleaching, the recovery images were acquired with 7 s interval in 10 min. The images and time-lapse movies were processed with ImageJ^46^.

### dsDNA-coupled cellulose beads pull-down assay

Proteins were incubated with dsDNA-coupled cellulose beads (Sigma) in the binding buffer (20 mM Tris-HCl pH 7.5, 150 mM NaCl), at 4°C for 1 h. After incubation, the beads were washed with the same binding buffer for three times. Then samples were eluted with buffer (20 mM Tris-HCl pH 7.5, 1 M NaCl). Input and eluted samples were analyzed by SDS-PAGE.

### Electromobility shift assays (EMSA)

100ng FAM labeled −20bp dsDNA was incubated with 50-0.2 μM AIM2 protein in 20 μL binding buffer (20 mM Tris-HCl pH 7.5, 150 mM NaCl) at 4°C for 30 min. After incubation, samples were resolved on a native 6% PAGE gel in 0.5× TBE buffer at 4°C. Gels were then analyzed by fluorescence imaging systems. The band intensity was measured through ImageJ and K_D_ was calculated by GraphPad Prism 8.0 software.

### Fluorescence resonance energy transfer (FRET) assay

ASC^PYD^ (106C) was labeled with sulfo-Cy3/Cy5 maleimide fluorophores in reaction buffer (20 mM HEPES pH 7.4, 150 mM NaCl, 1 mM TCEP). 1 μM AIM2 incubated with 0.5 μM DNA for 5 min before adding 2 μM ASC^PYD^ labeled with Cy3 and Cy5 fluorophores in reaction buffer (20 mM HEPES pH 7.4, 150 mM NaCl, 1 mM TCEP). Then immediately start recording the fluorescence signal. The data was processed and analyzed by GraphPad Prism 8.0 software.

### MBP pull-down assay

MBP-AIM2 was incubated with Dextrin Sepharose beads (GE Healthcare) in the binding buffer (150mM NaCl, 20 mM Tris-HCl pH7.5, 1 mM DTT) at 4°C for 1 h and washed three times with the binding buffer. Excessive VP22 protein was incubated with the protein-coated beads in the binding buffer at 4°C for 1 h. Then beads were washed three times with the binding buffer. The samples were eluted by elution buffer (150mM NaCl, 20 mM Tris-HCl pH7.5, 1 mM DTT, 20 mM maltose). Input and eluted samples were analyzed by SDS-PAGE.

### Plasmids construction for cellular assays

To generate the lentiviral vectors for doxycycline-inducible expression of AIM2 (pCW-GFP-AIM2), GFP-mAim2 and GFP-hAim2 fragments were amplified and inserted into pCW-Cas9-Blast vector (Addgene plasmid #83481). For the construction of transient expression plamids, Halo-mAim2, Flag-p202 and Vp22-sfgfp fragments were amplified and inserted into pCMV vector.

### Lentivirus packaging and Cell lines constructed

For lentivirus packaging, pCW-GFP-AIM2, NRF packaging plasmid and VSV-G envelope plasmid were co-transfected to HEK293T cells. The lentivirus was harvested from the supernatant at 48 h post-transfection and filtered with a 0.45 mm polyvinylidene fluoride filter. Then HEK293T cells were infected with the lentivirus. After 72 h post-transfection, cells were treated with 25 μg/ml doxycycline to induce the expression of GFP-AIM2 and then GFP-positive cells were sorted by Flow Cytometer. Cells were cultured and the expression of GFP-AIM2 was analyzed by western blot.

### Imaging AIM2-DNA puncta and perform FRAP assay in cells

The constructed HEK293T cells expressing GFP-AIM2 were plated onto collagen pre-coated coverslips. After 12 h post-plating, cells were induced by treating with 25 µg/mL doxycycline and transfected with 1 µg Cy5-labeled dsDNA oligoes for 5 h. Then imaging assay and FRAP assay were performed on a Zeiss LSM980 with Airyscan2 confocal laser scanning microscope equipped with a 63X /1.4 NA oil objective. The 488 nm laser were used for FRAP measurements. Bleaching was undertaken over a ∼1 mm radius using 100% laser power, and the post-bleaching images were collected from a z-stack of 5 µm every 8 s (mAIM2-DNA puncta) or 5 s (hAIM2-DNA puncta). Fluorescence intensity was measured using Software ZEN (Blue edition 3.8), and the resulted intensity traces were normalized according to the procedures in easyFRAP-web (https://easyfrap.vmnet.upatras.gr).

For imaging of AIM2 puncta in BMDMs or THP-1, BMDMs or THP-1 were transfected with Cy5-lableled 100-bp dsDNA for 8 h or BMDMs were infected with *F. novicida* at a MOI of 50 for 16 h. Then cells were imaged after immunofluorescence staining. Immunofluorescence images were captured and analyzed using Zeiss LSM980 with Airyscan2 confocal laser scanning microscope equipped with a 63X /1.4 NA oil objective.

For imaging of the inhibition of AIM2-ASC puncta formation by p202 and VP22, HEK293T cells were transfected with indicated plasmids by using lipo3000 for 12 h or 20h. Then cells were imaged after staining with 30 nM of HaloTag-JF549 dye (Promega) at 37°C for 30 min and immunofluorescence staining. The cells were imaged using the Multi-SIM imaging system (Beijing Nanolnsights-tech Co., Ltd) with a 100X/1.49 NA oil objective (Nikon CFI SR HP Apo).

### Immunofluorescence

The cells in immunofluorescence assays were fixed in 4% paraformaldehyde for 10 min, blocked with blocking buffer (3% normal donkey serum in PBST (PBS+0.1% Triton X-100) for 1 h, and incubated sequentially with primary antibodies including anti-AIM2 (1:200, Abcam, ab119791), anti-ASC (1:200, Santa Cruz Biotechnology, sc-271054), anti-Flag (1:200, Proteintech, 66008-4-Ig) diluted in blocking buffer at 4°C overnight. The specimen was washed twice with PBS and incubated with Alexa488/640 fluorochrome-conjugated secondary antibodies (1:1000, Invitrogen) diluted in blocking buffer for 1 h. Cells were then washed twice and finally imaged.

### Preparation of AIM2-DNA-ASC^PYD^ TEM grids

AIM2-DNA-ASC^PYD^ sample preparation is performed by mixing 5 μM AIM2, 1 μM 100-bp dsDNA and 0.2 mg/mL TEV in 150 mM NaCl buffer. After incubation for 15 min, 5 μM ASC^PYD^ was added to the reaction solution. After 10 min, sample was loaded onto the R1.2/1.3 Cu 300 mesh grids (Quantifoil) and then plunge-frozen into liquid ethane by using a Vitrobot (Thermo Fisher Scientific). The grids were stored in liquid nitrogen.

### Data collection and processing of cryo-ET

The prepared grids were subjected to a 300 kV Titan Krios microscope (FEI, Thermo Fisher Scientific), equipped with a post-column energy filter (Gatan) and a K2 Summit direct detector camera (Gatan). Using SerialEM software^47^, tilt-series were acquired from −50° to +50°, with 2° increments. Images were recorded in movie mode at 11 frames per second, with an object pixel size of 1.36 Å (magnification of 105,000x) and a defocus of −3 to −6 μm with a dose-symmetric tilt scheme using the beam-shift method^48^. The dose was set at 2.5 e/Å2 per tilt image collected over frames.

Raw images of the tilt series were processed in Warp for motion correction^49^, contrast transfer function (CTF) estimation and tilt series sorting. Tilt series was aligned using the patch-tracking method included in the IMOD software package^50^. Then the alignment parameters were sent back to Warp, and the tomograms were reconstructed at 4× binning (voxel size = 5.44 Å).

For data processing, motion correction, CTF estimation, dose filtering and tilt-series sorting were performed in Warp. The dataset was processed with the denoising network in Warp for visualization purposes only.

### Data collection and processing of cryo-EM

Data collection of cryo-EM was performed on a 300 kV Titan Krios electron microscope (FEI) equipped with K2. Summit camera (Gatan) and a GIF Quantum energy filter operated with a slit width of 20 eV. All cryo-EM super-resolution micrographs were collected automatically using the Serial-EM package, yielding an image stack with a pixel size of 0.52 Å. Images were recorded at a defocus range of −1.2 μm to −1.8 μm, with a total electron dose of 60 e/Å2 over 32 movie frames. Images were recorded by beam-image shift data collection methods^47^. Each movie stack was motion-corrected by MotionCor2^51^. The dose-weighted micrographs were kept for further image processing using CryoSPARCv4.5.1^52^.

The exact defocus value and contrast transfer function (CTF) of each micrograph were estimated using CryoSPARC’s patch CTF estimation tool. Particles were automatically picked using Blob picking and Templet picking. For the data of ASC^PYD^ filament, a total of 402,816 particles were auto-picked from 581 micrographs. After 2D classification, 81,574 particles with good features were kept for further data processing. A total of 45,400 particles were used for further homogeneous refinement and non-uniform refinement with C3 symmetry resulting in a 2.7 Å map, whose resolution was estimated based on the gold-standard Fourier shell correlation (FSC) with 0.143 criterion.

For the atomic model of ASC^PYD^ filament, use the structure of ASC^PYD^ filament predicted by AlphaFold-Multer^53^, as the initial model, was fitted into the cryo-EM map using UCSF Chimera^54^. The resulting model was then manually rebuilt in COOT^55^ and further refined by real space refinement in PHENIX^56^. Structural figures were generated using PyMOL^57^ and ChimeraX^58^.

### BMDMs stimulation and infection

BMDMs were plated onto 12-well plates at a density of 1 × 10^6^ cells per well and incubated overnight before use. Stimulation and infection assays were performed as described previously.^59^ For stimulation with poly(dA:dT), each well of BMDMs was transfected with 2 μg poly(dA:dT) by using Xfect reagent for 5 h. For *F. novicida* infection, *F. novicida* was grown in TSB supplemented with 0.2% L-cysteine overnight at 37 °C. Each well of BMDMs was infected with *F. novicida* at a MOI of 50 for 16 h. For cell death analysis, each well of BMDMs was added with 20 nM SYTOX and imaged by confocal at indicated time point.

### *F. novicida* infected mice

6- to 8-week-old *Aim2^WT^ mice*, *Aim2^IDR_M^*mice and *Aim2^HIN_M^* mice were used for infections. Mice were infected subcutaneously with 5 × 10^5^ CFU of *F. novicida* in 200 μl PBS. For survival analysis, infected mice were continuously observed for 12 days. To evaluate the bacterial load, the spleen, liver and lung tissues of infected mice were collected and homogenized to perform plaque assays.

### Experimental colitis and disease assessment

Mice were administered DSS (3% w/vol; molecular weight: 36-50 kDa; MP Biomedicals) in drinking water for 7 days, followed by regular drinking water for 3 days. Clinical signs of colitis were evaluated based on the DAI score including body weight loss, occult blood, and stool consistency. In brief, the weight loss score was determined as follows: 0, no weight loss; 1, loss of 1-5% original weight; 2, loss of 6-10% original weight; 3, loss of 11-20% original weight; 4, loss of >20% original weight. The bleeding score was determined as follows: 0, no blood by using Haemoccult analysis; 1, positive Haemoccult; 2, visible blood traces in stool; and 3, gross rectal bleeding. The stool score was graded as follows: 0, well-formed pellets; 1, semiformed stools that did not adhere to the anus; 2, pasty semiformed stool that attached to the anus; and 3, liquid stools that attached to the anus.

For the histological activity index evaluation, harvested colon sections were stained with H&E. The histological activity index was assessed according to the loss of goblet cells and crypts and the extent of inflammatory cell infiltration. Crypt gland destruction was scored as follows: 0, normal; 1, a small amount of crypt loss; 2, a large amount of crypt loss; 3, extensive crypt loss. Goblet cells loss was scored as follows: 0, normal; 1, a small amount of goblet cells loss; 2, a large amount of goblet cells loss; 3, extensive goblet cells loss. Inflammatory cell infiltration was scored as follows: 0, normal; 1, infiltration around crypt glands; 2, infiltration of mucosal muscle layer; 3, extensive infiltration of the mucosal muscle layer with oedema; 4, infiltration of the submucosal layer.

### Immunoblot and ELISA analysis

For caspase analysis, each well of BMDMs (contain 400 μl supernatant) was added with 100 μl 5× SDS loading buffer (15 g tris, 74 g glycine, 5 g SDS, 1× protease inhibitors, 1× phosphatase inhibitors in 1L ddH2O). Then combined supernatant and protein lysates were collected. For other analysis, the supernatants were removed and cells were washed with PBS, then each well of BMDMs was added with 80 μl RIPA lysis buffer (Solarbio) and collected. Collected samples were denatured at 95 °C for 10 min and then analyzed by gel electrophoresis. Protein was then wet transferred to a 0.45 mm PVDF membrane (Millipore) in ice-cold transfer buffer (25mM Tris, 192mM glycine, 20% methanol). The membrane was blocked with 5% skim milk in TBST for 1 h and then incubated with indicated primary antibodies overnight at 4°C : anti-AIM2 rabbit pAb (ABclonal, A25252,1:1000), anti-ASC/TMS1 rabbit mAb (CST, 17507, 1:1000), anti-caspase-1 mouse mAb (AdipoGen, AG-20B-0042-C100, 1:1000), anti-GSDMD rabbit mAb (Abcam, ab209845, 1:1000), anti-cleaved caspase-8 rabbit mAb (CST, 8592, 1:1000), anti-caspase-3 rabbit mAb (CST, 14220, 1:1000), anti-cleaved caspase-3 rabbit pAb antibody (CST, 9661, 1:1000), anti-pMLKL rabbit mAb (CST, 37333, 1:1000), anti-pRIPK3 rabbit mAb (CST, 91702, 1:1000), anti-MLKL rabbit mAb (ABclonal, A21894), anti-RIPK3 rabbit mAb (CST, 95702, 1:1000), anti-β-actin mouse mAb (ABclonal, WH278304). After being incubated with primary antibody and washed with TBST, the membrane was incubated with appropriated second antibodies for 1 h: HRP-conjugated goat anti-rabbit IgG (ABclonal, AS014, 1:6000), anti-mouse IgG (ABclonal, AS003, 1:6000). After washing with TBST, sample were visualized by using SuperSignal West Atto Substrate and imaged by Tanon-5200 machine (Tanon).

For ELISA analysis, cell supernatant was collected and analyzed by ELISA kit (Beyotime, PI553) according to the manufacturer’s instructions.

### Statistical analysis

GraphPad Prism 8.0 software was used to perform statistical analysis. Statistical significance was determined by two-tailed Student’s t test or one-way ANOVA. Detailed information is available in the figure legends.

---

**Figure S1.**
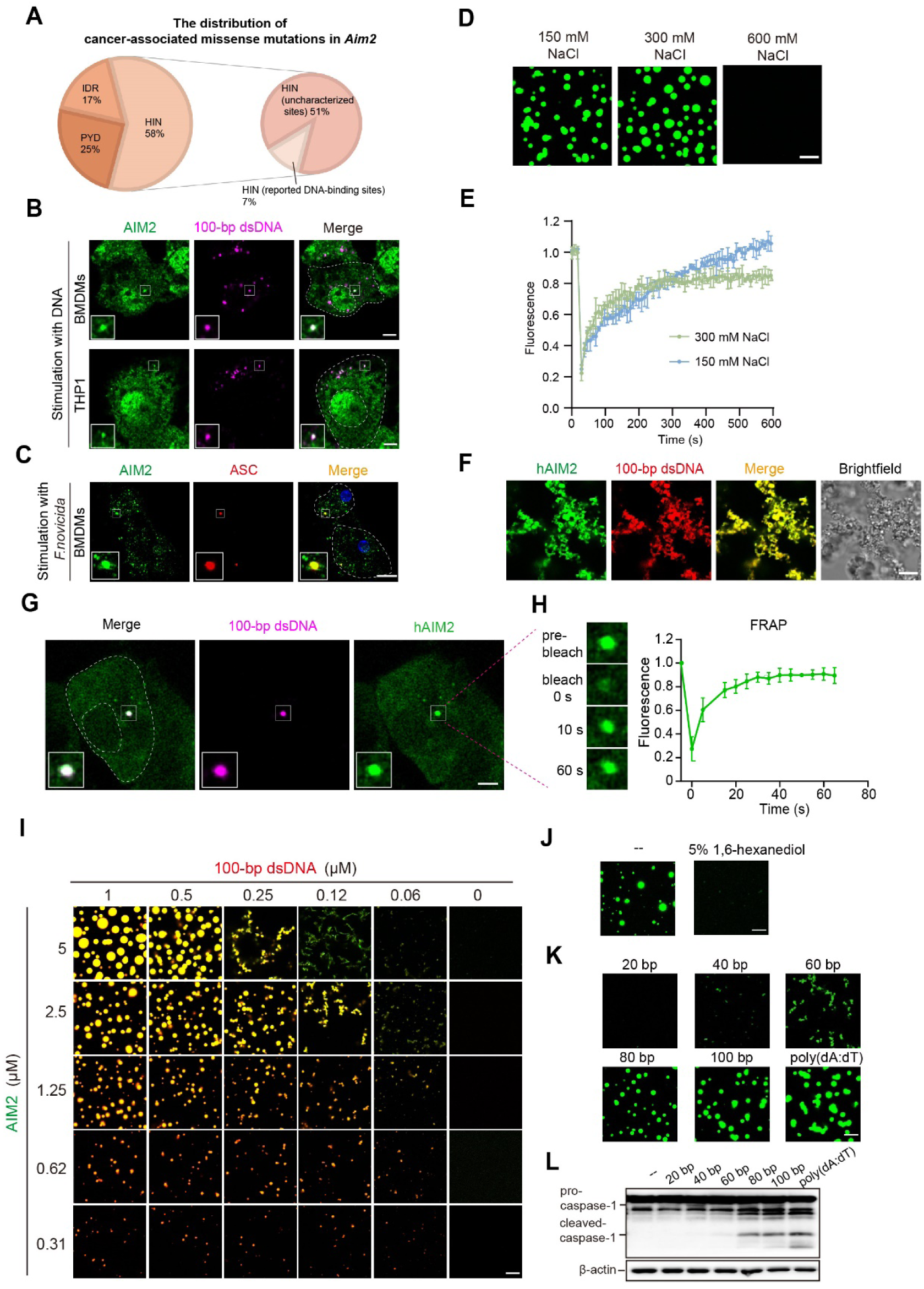
dsDNA induces AIM2 to form liquid-phase condensation *in vitro* and in cells. (A) The distribution of cancer-associated missense mutations in *Aim2*. Data are from the Cancer Genome Atlas (TCGA) database. (B) Immunofluorescence images of BMDMs and THP-1 at 8 h after transfecting with cy5-labeled 100-bp dsDNA. The AIM2-DNA puncta highlighted by the white box is magnified at the bottom left. Scale bar, 4 μm. 3 μm for magnified images. (C) Immunofluorescence images of BMDMs at 16 h after *F. novicida* infection. The AIM2-ASC puncta highlighted by the white box is magnified at the bottom left. Scale bar, 10 μm. 4 μm for magnified images. (D) Images of the condensates formed by 5 μM AF488-labeled mAIM2 and 1 μM 100-bp dsDNA in indicated salt concentration. Scale bar, 10 μm. (E) FRAP analysis of mAIM2-DNA droplets in 150 mM NaCl buffer or 300 mM NaCl buffer. These droplets were formed by incubating 5 μM AIM2 with 1 μM DNA. Values shown are means ± SD. n=3 condensates. (F) Images of the condensates formed by 5 μM AF488-labeled hAIM2 and 1 μM cy3-labeled 100-bp dsDNA in 300 mM NaCl buffer. Scale bar, 10 μm. (G) Representative live-cell images of hAIM2-dsDNA puncta in HEK293T cells expressing GFP-hAIM2 (Tet-on) at 5h after transfecting with 1 μg/mL cy5-labeled 100-bp dsDNA and treating 25 µg/mL doxycycline to induce the expression of GFP-hAIM2. The hAIM2-DNA puncta highlighted by the white box is magnified at the bottom left. scale bar, 4 μm; 3 μm for magnified images. (H) FRAP analysis of hAIM2-DNA puncta that shown in (G). Left panel: time-lapse images of hAIM2-DNA puncta before and after photobleaching. 3 μm for magnified images. Right panel: quantification of FRAP data. Values shown are means ± SD. n=3 puncta. (I) Phase separation diagram of AF488-labeled mAIM2 and cy3-labeled 100-bp dsDNA in 300 mM NaCl buffer. Merged images are shown. Scale bar, 10 μm. (J) Images of the condensates formed by 5 μM AF488-labeled mAIM2 and 1 μM DNA in 300 mM NaCl buffer after adding buffer (left panel) or 5% 1,6-hexanediol (right panel). (K) Images of the condensates formed by 5 μM AF488-labeled mAIM2 and 1 μM indicated length DNA in 300 mM NaCl buffer. Scale bar, 10 μm. (L) Immunoblot analysis of pro- and cleaved-caspase-1 in BMDMs at 5 h after stimulation with 2 μg indicated length DNA.

**Figure S2.**
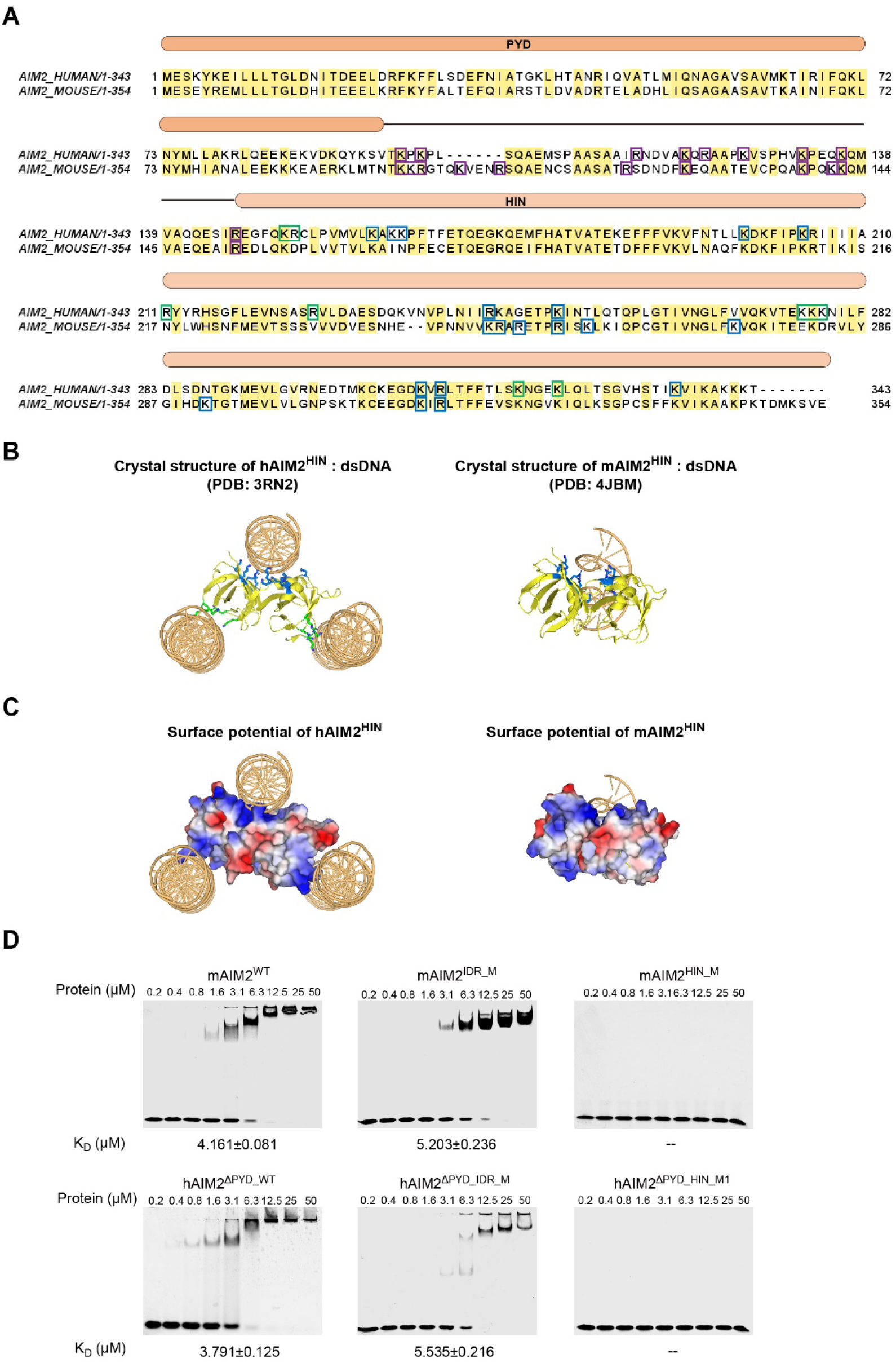
The analysis of amino acid sequences, protein structures, surface potential and mutants of AIM2. (A) Sequences of hAIM2 and mAIM2 were aligned and analyzed by Jalview. Domain elements of AIM2 are indicated above the sequence alignment. Conserved residues are shaded in yellow. Positively charged residues within IDR are in purple boxes. Main DNA binding sites are in blue boxes and additional DNA binding sites are in green boxes. (B) Left panel: crystal structure of hAIM2^HIN^:dsDNA (PDB: 3RN2). Right panel: crystal structure of mAIM2^HIN^:dsDNA (PDB: 4JBM). Main DNA binding sites are shown in blue stick. Additional DNA binding sites are shown in green stick. (C) Left panel: surface potential of hAIM2^HIN^. Right panel: surface potential of mAIM2^HIN^. Positive charge is represented by blue and negative charge is represented by red. (D) Electrophoretic mobility shift assay (EMSA) of mAIM2WT and its mutants (top panel), hAIM2 ΔPYD_WT and its mutants (bottom panel). KD shown are means ± SD, n=3.

**Figure S3.**
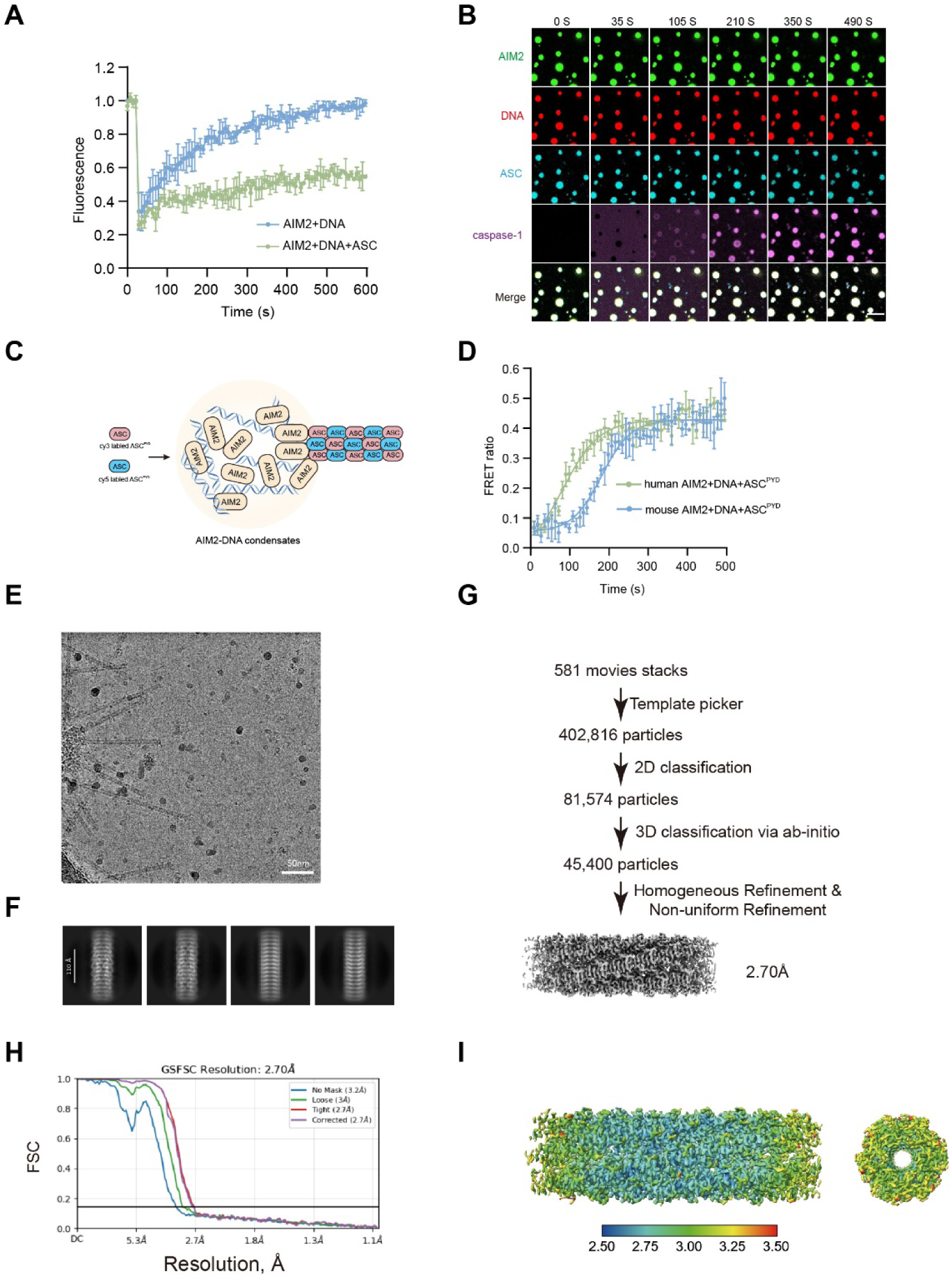
AIM2-DNA condensation facilitates inflammasome and PANoptosome assembly. (A) FRAP analysis of AIM2–DNA condensates after the addition of ASC. (B) Time-lapse images of droplets that formed as described in Figure 3A. Scale bar, 5 μm. (C) Schematic diagram of FRET assay. ASC^PYD^ labeled with Cy3 and Cy5 fluorophores incubated with AIM2-DNA condensates. (D) FRET assay of AIM2-DNA condensates nucleate ASC^PYD^ filament formation. 1 μM AIM2 incubated with 0.5 μM DNA for 5 min before adding 2 μM ASC^PYD^ labeled with Cy3 and Cy5 fluorophores. Then immediately start recording the fluorescence signal. (E) Representative raw micrograph of ASC^PYD^ filament from AIM2-DNA-ASC^PYD^. (F) Selected 2D class averages of ASC^PYD^ filament. (G) Data processing flow chart. (H) Gold-standard FSC curves used for global-resolution estimates within cryoSPARC. (I) Local resolution distribution of ASC^PYD^ filament.

**Figure S4.**
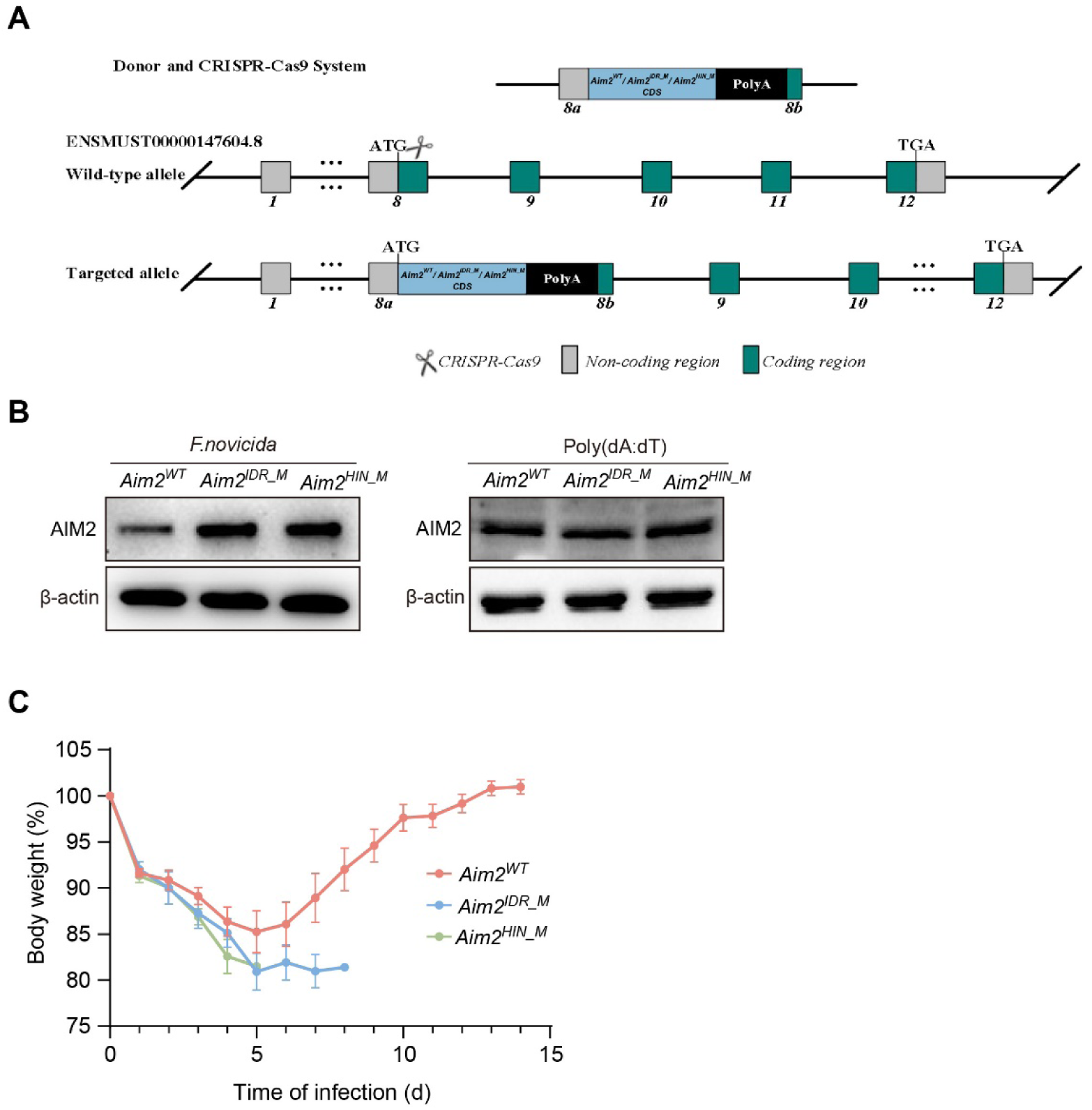
The generation of *Aim2^WT^*, *Aim2^IDR_M^*, *Aim2^HIN_M^* mice and infection assay. (A) Schematic of generate three types of transgenic mice (*Aim2^WT^*, *Aim2^IDR_M^*and *Aim2^HIN_M^* mice) by CRISPR-Cas9. (B) Immunoblot analysis of AIM2 expression in *Aim2^WT^*, *Aim2^IDR_M^* and *Aim2^HIN_M^*BMDMs after stimulation with *F. novicida* (left panel) or poly(dA:dT) (right panel). (C) Weight loss of *Aim2^WT^*, *Aim2^IDR_M^* and *Aim2^HIN_M^*mice after infection with 5×10^5^ CFU of *F. novicida*.

**Figure S5.**
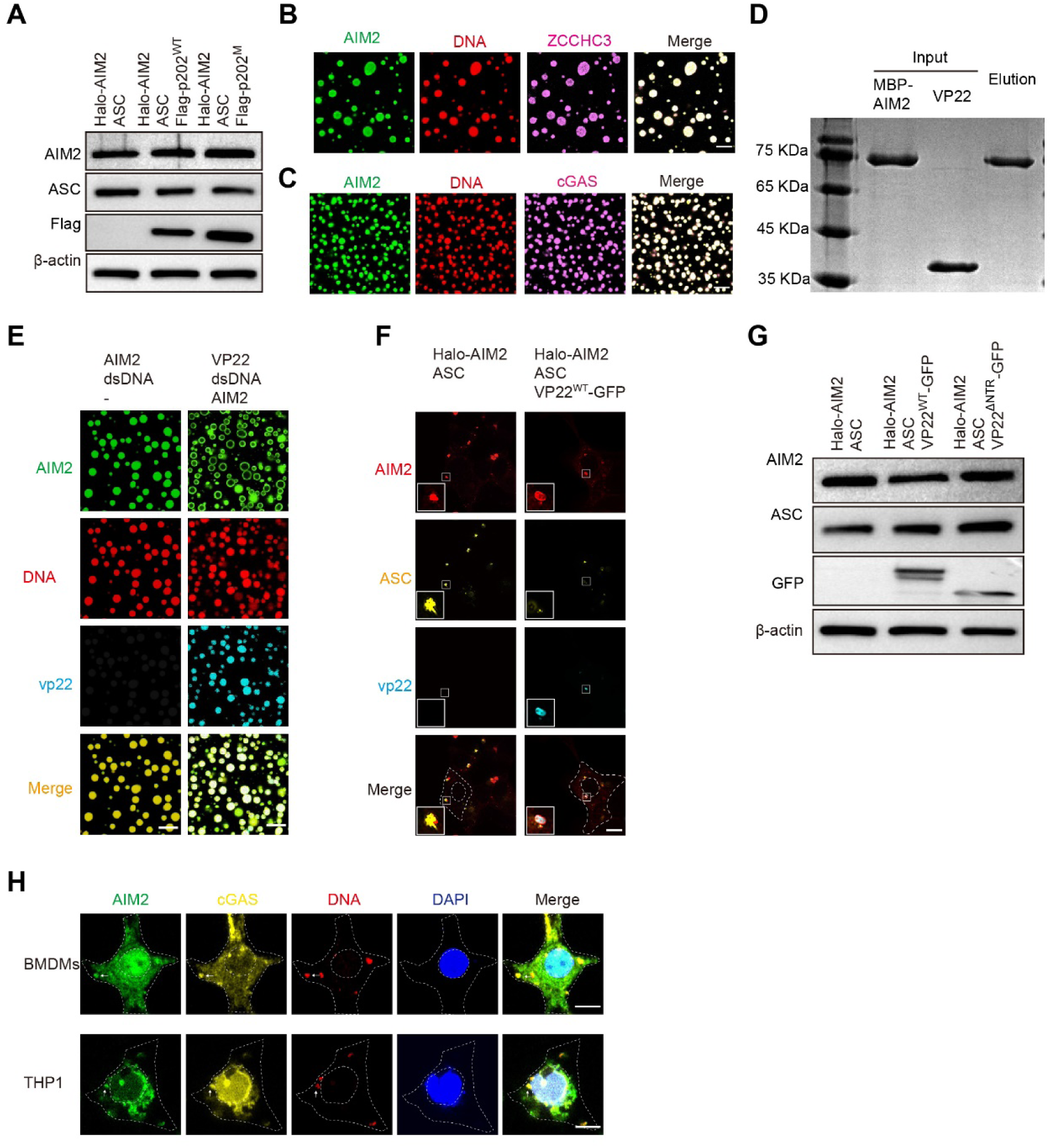
Host- and pathogen-mediated regulation of AIM2-DNA phase separation. (A) Immunoblot analysis of the expression of AIM2, ASC, p202 in Figure 6B. (B) Images of droplets formed after co-incubating 5 μM AF488-labeled AIM2, 10 μM AF647-labeled ZCCHC3 and 1 μM cy3-100bp dsDNA. (C) Images of droplets formed after co-incubating 5 μM AF488-labeled AIM2, 10 μM AF647-labeled cGAS and 1 μM cy3-100bp dsDNA. (D) MBP pull-down assay of MBP-AIM2 and VP22. (E) Images of 5 μM AF488-labeled AIM2 incubated with 1 μM cy3-labeled 100-bp dsDNA (left panel) or preformed VP22-DNA condensates that is formed by 10 μM AF647-labeled VP22WT and 1 μM cy3-labeled 100-bp dsDNA (right panel). Scale bar, 10 μm. (F) Immunofluorescence images of HEK293T cells transfected with indicated plasmids for 24 h. Scale bar, 10 μm. 5 μm for magnified images. (G) Immunoblot analysis of the expression of AIM2, ASC, VP22 in Figure 6E. (H) Immunofluorescence images of BMDMs and THP-1 at 8 h after transfecting with cy5-labeled 100-bp dsDNA. Arrowheads indicate AIM2-cGAS-DNA puncta. Scale bar, 5 μm.

**Video S1. Z-axis fly-through of the reconstructed tomogram showing the intact AIM2-DNA condensates. Arrowheads indicate ASC^PYD^ filaments.**

**Table S1.** Protein truncations.

| Truncation Name |  | Amino acid Sequence |
| --- | --- | --- |
| mAIM2 | ΔPYD | T96-E354 |
|  | ΔIDR-HIN | M1-N95 |
|  | ΔHIN | M1-R152 |
|  | ΔPYD-IDR | E153-E354 |
|  | mAIM2-FUS | FUS<sup>IDR</sup>(G156-D212) replaces mAIM2<sup>IDR</sup>(T96-R152) |
|  | mAIM2-cGAS | cGAS<sup>IDR</sup>(T72-R128) replaces mAIM2<sup>IDR</sup>(T96-R152) |
| hAIM2 | ΔPYD | T96-T343 |
|  | ΔIDR-HIN | M1-V95 |
|  | ΔHIN | M1-R146 |
|  | ΔPYD-IDR | E147-T343 |
| mRIPK1<sup>RHIM-DD</sup> |  | S510-S656 |
| mRIPK3<sup>RHIM</sup> |  | T405-K486 |
| mcaspase-8<sup>DED</sup> |  | M1-S188 |
| VP22<sup>ΔNTR</sup> |  | R174-E301 |

**Table S2.** Protein Mutations.

| Mutation Name | Mutation sites |
| --- | --- |
| mAIM2<sup>IDR_M</sup> | K97D, K98D, R99D, K103D, R107D, R120D, K126D, K138D, K141D, K142D, R152D |
| mAIM2<sup>HIN_M</sup> | K248D, R249D, R251D, R255D, K258D |
| mAIM2<sup>ΔPYD+Sites</sup> | D158R, N217R, V232R, E280K, D282K |
| mAIM2<sup>PYD_M</sup> | D15R, Y74R |
| hAIM2<sup>ΔPYD_IDR_M</sup> | K97D, K99D, R115D, K120D, R122D, K126D, K132D, K136D, R146D |
| hAIM2<sup>ΔPYD_HIN_M1</sup> | K160D, K162D, K163D, K198D, K204D, R244D, K251D, K309D, R311D, K335D |
| hAIM2<sup>ΔPYD_HIN_M2</sup> | K151D, K152D, R211D, R226D, K276D, K277D, K278D, K319D, K323D |
| hAIM2<sup>PYD_M</sup> | D15R, Y74R |
| p202<sup>M</sup> | K48A, R224A |
| mZBP1<sup>mut</sup> | K42A, K43A, K119A |

**Table S3.** Cryo-EM data collection, refinement and validation statistics.

|  | ASC<sup>PyD</sup> Filament<br>(PDB 9U8K) (EMD-63955) |
| --- | --- |
| **Data collection processing** |  |
| Magnification | 130,000 |
| Voltage (kV) | 300 |
| Electron exposure (e<sup>-</sup>/Å<sup>2</sup>) | 60 |
| Defocus rang (μm) | -1.2 ~ -1.8 |
| Pixel size (Å) | 0.52 |
| Symmetry imposed | C3 |
| Initial particle images (no.) | 81,574 |
| Final particle images (no.) | 45,400 |
| Map resolution (Å) | 2.70 |
| FSC threshold | 0.143 |
| Map resolution range (Å) | - |
| **Refinement** |  |
| Initial model used (PDB code) | None |
| Model resolution (Å) | 3.13 |
| FSC threshold | 0.5 |
| Model resolution range (Å) | - |
| Map sharpening *B* factor (Å) | -75.5 |
| Model composition |  |
| Non-hydrogen atoms | 27482 |
| Protein residues | 3465 |
| *B* factor (Å<sup>2</sup>) |  |
| Protein | 70.51 |
| R.m.s. deviations |  |
| Bond lengths (Å) | 0.005 |
| Bond angles (°) | 0.690 |
| Validation |  |
| MolProbity score | 1.56 |
| Clashscore | 7.82 |
| Poor rotamers (%) | 0 |
| Ramachandran plot |  |
| Favored (%) | 97.31 |
| Allowed (%) | 2.69 |
| Disallowed (%) | 0 |

